# Single-cell RNA profiling of *Plasmodium vivax*-infected hepatocytes reveals parasite- and host- specific transcriptomic signatures and therapeutic targets

**DOI:** 10.1101/2022.02.01.478648

**Authors:** Anthony A Ruberto, Steven P Maher, Amélie Vantaux, Chester J Joyner, Caitlin Bourke, Balu Balan, Aaron Jex, Ivo Mueller, Benoit Witkowski, Dennis E Kyle

## Abstract

The resilience of *Plasmodium vivax*, the most widely distributed malaria-causing parasite in humans, is attributed to its ability to produce dormant liver forms known as hypnozoites, which can activate weeks, months, or even years after an initial mosquito bite. The factors underlying hypnozoite formation and activation are poorly understood, as is the parasite’s influence on the host hepatocyte. Here, we shed light on transcriptome-wide signatures of both the parasite and the infected host cell by sequencing over 1,000 *P. vivax*-infected hepatocytes at single-cell resolution. We distinguish between replicating schizonts and hypnozoites at the transcriptional level, identifying key differences in transcripts encoding for RNA-binding proteins associated with cell fate. In infected hepatocytes, we show that genes associated with energy metabolism and antioxidant stress response were upregulated, and those involved in the host immune response downregulated, suggesting both schizonts and hypnozoites alter the host intracellular environment. The transcriptional markers in schizonts, hypnozoites, and infected hepatocytes revealed here pinpoint potential factors underlying dormancy and can inform therapeutic targets against *P. vivax* liver-stage infection.

## INTRODUCTION

Malaria is a significant global health burden, with an estimated 421 million cases and 627,000 deaths per year (World Health Organization, 2021). At least five *Plasmodium* species are known to infect humans, with *P. vivax* responsible for the majority of cases outside Africa (World Health Organization, 2021). All *Plasmodium* parasites that infect humans have an obligatory developmental stage in the liver where the parasite undergoes asexual reproduction known as schizogony within a hepatocyte before releasing thousands of merozoites into the blood. In *P. vivax*, a proportion of parasites known as hypnozoites forgo immediate division in hepatocytes, and persist in the liver for weeks, months, or years before activating to initiate schizogony resulting in a relapse blood infection (Adams and Mueller, 2017; Krotoski et al., 1982). Hypnozoites are thought to cause up to 90% of cases in certain geographical regions (Adekunle et al., 2015; Commons et al., 2020; Robinson et al., 2015), and mathematical models predict that eliminating *P. vivax* malaria will be difficult without interventions that target hypnozoite reservoirs (White et al., 2018).

The factors influencing the biogenesis, persistence, and activation of hypnozoites are poorly understood. Studies have found that hypnozoites in temperate versus tropical regions have long and short latencies, respectively, suggesting a correlation between geographical region and relapse patterns (Battle et al., 2014; Huldén et al., 2008; Krotoski et al., 1986; White, 2011). These findings suggest the developmental trajectory is genetically encoded, and support the hypothesis that it is somehow pre-programmed in the sporozoite (Lysenko et al., 1977; Ungureanu et al., 1976); but other contributing factors from the host could play a role in determining whether the sporozoite becomes a schizont or hypnozoite (Schäfer et al., 2021).

Our understanding of *P. vivax* liver stage biology at the cellular and molecular level is in its nascency for two overarching reasons. First, protocols to cryopreserve and continuously maintain erythrocytic stage parasites *in vitro* are lacking, necessitating the collection of gametocyte-infected blood from human volunteers in *P. vivax* malaria-endemic regions to generate the sporozoites required to initiate an infection. New laboratory models have recently been developed which can partially alleviate the need to obtain gametocytes from human-infected blood (Mikolajczak et al., 2015; Schäfer et al., 2020), yet these sources are expensive and each have distinct logistical hurldes. In addition to the sporozoite limitation, upon initiating a *P.* vivax-infected hepatocyte culture, the high ratio of noninfected versus infected hepatocytes has historically made it difficult to isolate and study these forms in sufficient numbers. Robust *in vitro* pre-erythrocytic platforms capable of long-term culture of sufficient numbers of *P. vivax* liver stages have only recently been developed (Maher et al., 2021; Roth, Maher et al., 2018). For these reasons, much of our knowledge of hypnozoite biology at the molecular level remains limited to studies using *P. cynomolgi*—a parasite capable of causing relapsing infections in non-human primates for which molecular tools are available (Chua et al., 2019; Cubi et al., 2017; Voorberg-van der Wel et al., 2017, 2020). However, it is unknown whether mechanisms regulating hypnozoite formation and dormancy are consistent across the different relapsing malaria parasites.

Transcriptome-wide profiling of *P. vivax* liver stages has previously been performed using a bulk RNA sequencing strategy (Gural et al., 2018). While informative, the data provide insights only into average transcript levels derived from either mixed (schizont and hypnozoite) or hypnozoite-only populations, masking any transcriptional variation that might exist between individual parasites. Second, the parasite’s potential impact on the hepatocyte is not considered, barring analysis of host-pathogen interactions. Nevertheless, these shortcomings highlight a need to better understand differences between individual parasites, as well as how the parasite alters the hepatocyte. Advancements in these areas could help elucidate parasite- and host-factors important for the parasite’s liver stage development.

Single-cell RNA-sequencing (scRNA-seq) methods offer new ways of studying cell-to-cell variation (Aldridge and Teichmann, 2020), and have already allowed for novel insights into host-pathogen interactions in various disease contexts (Penaranda and Hung, 2019). In *Plasmodium* spp., scRNA-seq analyses have so far revealed that individual parasite forms exist on a spectrum of transcriptomic states (Bogale et al., 2021; Howick et al., 2019; Poran et al., 2017; Real et al., 2021; Reid et al., 2018; Ruberto et al., 2021a), and applying them to study the biology of *P. vivax* is a burgeoning area of research (Mancio-Silva et al., 2022; Ruberto et al., 2021b; Sà et al., 2020). Droplet-based techniques, such as 10x Genomics’ gene expression platform (Zheng et al., 2017), allow for the sequencing of several thousands of cells, increasing the chances of observing rare cell populations (e.g. hepatocytes infected with *P. vivax*). We therefore presumed that applying scRNA-seq to *P. vivax* liver forms derived using an *in vitro* platform would prove a fruitful avenue of inquiry into the parasite’s biology at the transcription level.

Using scRNA-seq, we characterize transcriptomic signatures of parasite and host in an *in vitro P. vivax* liver stage model. We find differences in gene expression between replicating schizonts and hypnozoites, and reveal variation, previously unobservable by bulk sequencing approaches, between individual hypnozoites. In infected hepatocytes, we find that genes associated with energy metabolism and antioxidant stress response are upregulated, and those involved in the host immune response downregulated. Various host genes associated with these processes are found to be exclusively upregulated during hypnozoite infection. Our study elucidates the transcriptional signatures among infected hepatocytes and presents potential mechanisms *P. vivax* can use to either exploit its host for rapid proliferation or to quietly persist, remaining undetected by the host immune system.

## RESULTS

### Strategy used to study *P. vivax* liver stage biology

Primary human hepatocytes were cultured and infected with *P. vivax* sporozoites on a 384-well plate platform (Maher et al., 2021; Roth, Maher et al., 2018). Figure 1A outlines the workflow for the experiment. Following infection with sporozoites, cultures were assessed at two endpoints: one containing both schizonts and hypnozoites at day 5 post-infection, and the other containing solely hypnozoites at day 9 post-infection. Schizonts were eliminated from day 9 post-infection cultures by adding the phosphatidylinositol 4-kinase (PI4K) inhibitor MMV390048 (1 µM) to the culture media on days 5, 6 and 7 post-infection (Gural et al., 2018; Paquet et al., 2017). For each collection time point, we simultaneously initiated two sets of hepatocyte cultures: one infected with sporozoites for quantification of immunofluorescent-labeled parasites by high-content imaging (HCI), and one infected with sporozoites for transcriptional profiling using a high-throughput, droplet-based scRNA-seq workflow (Zheng et al., 2017). As controls, a third set of naïve cultures were left uninfected and processed for scRNA-seq alongside cultures at days 5 and day 9 post-infection. At each collection point, a target of approximately 10,000 individual hepatocytes were partitioned into droplets prior to the generation of RNA libraries. Droplets contained primers with unique nucleotide sequences (cellular barcodes) that were incorporated into the transcript sequence during the reverse transcription step, allowing for the identification of each transcript’s cellular origin. After sequencing the single-cell libraries, we mapped the resulting reads onto a customized reference transcriptome containing human (Ensembl, v103) and *P. vivax* PVP01 (PlasmoDB, v51) information. The *P. vivax* PVP01 transcript information included the nucleotides encoding for the gene-flanking untranslated regions (UTRs) (Siegel, Chappell et al., 2020), which is important because of the poly-dT capture strategy of the technology used in our workflow. This information partially alleviates the challenges associated with transcription quantification (Ruberto et al., 2021b; Sà et al., 2020). Post mapping, we used the transcript information (i.e. human or parasite encoding) with their designated cellular barcodes to assess the number of transcripts in each cell, and to determine which hepatocytes were infected.

**Figure 1.**
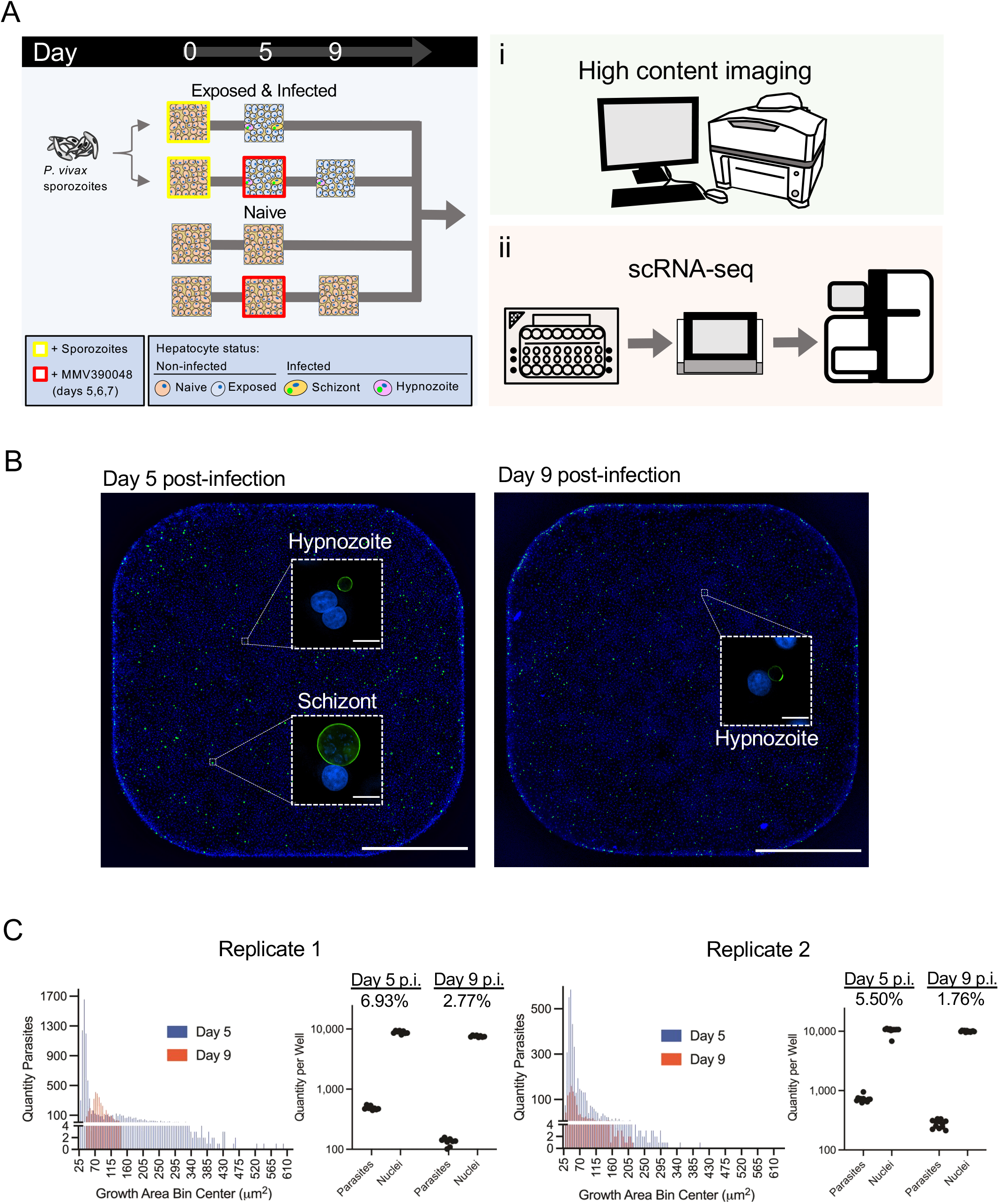
Design and validation of detection of P. vivax liver stages using high-content imaging and scRNA-seq. (A) Schematic illustrating the sample preparation and data generation pipeline used to assess P. vivax liver stage infection rates and gene expression. (B) Representative high content images of primary hepatocytes infected with *P. vivax* on day 5 (left) and day 9 (right) post-infection (p.i). Images were captured from one well from a 384-well plate with a 4x objective. Inset: one field of view (white dashed box) from the same well captured with a 20x objective. Cells were stained with DAPI (blue) and *Pv*UIS4 (green). White bar represents 1mm, inset bars represent 10µm. Images are representative of replicate 1. (C) Histogram displaying the distribution of parasite growth areas fixed at day 5 or day 9 post-infection (left) and net quantity of parasites and hepatocyte nuclei (right). Net infection rate is indicated for each endpoint. Growth and infection metrics are shown for both replicates.

As expected, we detected hepatocytes infected with either form of the parasite at 5 days post-infection, and only hypnozoites at 9 days post-infection following exposure to MMV390048 (Figure 1B **and Figure 1—figure supplement 1**). Visual assessment of infected cultures fixed and stained with an antibody against Upregulated in Infectious Sporozoites 4 (*Pv*UIS4), which localizes to the parasite’s parasitophorous vacuole membrane, confirmed that the parasites we subjected to scRNA-seq had properly invaded hepatocytes and initiated development into hypnozoites and schizonts. Furthemore, hypnozoites displayed typical morphology relative to those found in humanized mice (Mikolajczak et al., 2015) and *in vitro* (Roth, Maher et al., 2018). Figure 1C displays the distribution of individual parasite growth areas, number of parasites detected, and parasitemia levels at each endpoint. Across replicates, the parasitemia ranged from 5.50-6.93% at 5 days post-infection and 1.76-2.77% at 9 days post-infection. Together, the high infection rates coupled with the high-throughput capacity of 10x Genomics’ single-cell isolation and sequencing workflow allowed for sufficient material to be generated for successful profiling of both parasite and host transcriptomes.

### *P. vivax* schizonts and hypnozoites are distinguishable at single-cell resolution

We first analyzed the reads mapping to *P. vivax* transcripts to identify schizonts and hypnozoites, as well as to assess the extent to which these forms differed at single-cell resolution. In total, we characterized 1,438 parasite transcriptomes across two independent experimental replicates (**Figure 2— figure supplement 1A**). The number of parasite transcriptomes profiled corroborated the parasitemia levels quantified using high-content imaging. This correlative evidence suggests that our cell and gene filtering cutoffs effectively eliminated problematic cells (i.e. poorly captured, dying, or dead) (**Figure 2—figure supplement 1B and Figure 2—source data 1**). Low dimensional representation of the data revealed two populations of parasites in samples profiled at 5 days post-infection (Figure 2A**, left**). In contrast, and as expected, samples profiled at 9 days post-infection contained one population (Figure 2A**, right**). Given that the population of cells profiled at day 9 post-infection, from which schizonts were chemically removed, overlapped with one of the two cell populations profiled at 5 days post-infection, we deduced that the non-overlapping population of cells encoded for schizonts. Consistent with our reasoning, an unsupervised clustering algorithm (Traag et al., 2019) unbiasedly assigned the cells to one of the two clusters (Figure 2B **and Figure 2—figure supplement 1C**). As expected, the number of transcripts detected, the unique molecular identifier (UMI) counts (a readout for absolute transcript abundances (Kivioja et al., 2011)), and the transcript levels of the early liver stage development marker liver stage-specific protein, LISP2 (PVP01_0304700) (Gupta et al., 2019), all displayed greater quantities in the schizont cluster (C2) relative to the hypnozoite cluster (C1) (Figure 2C).

**Figure 2.**
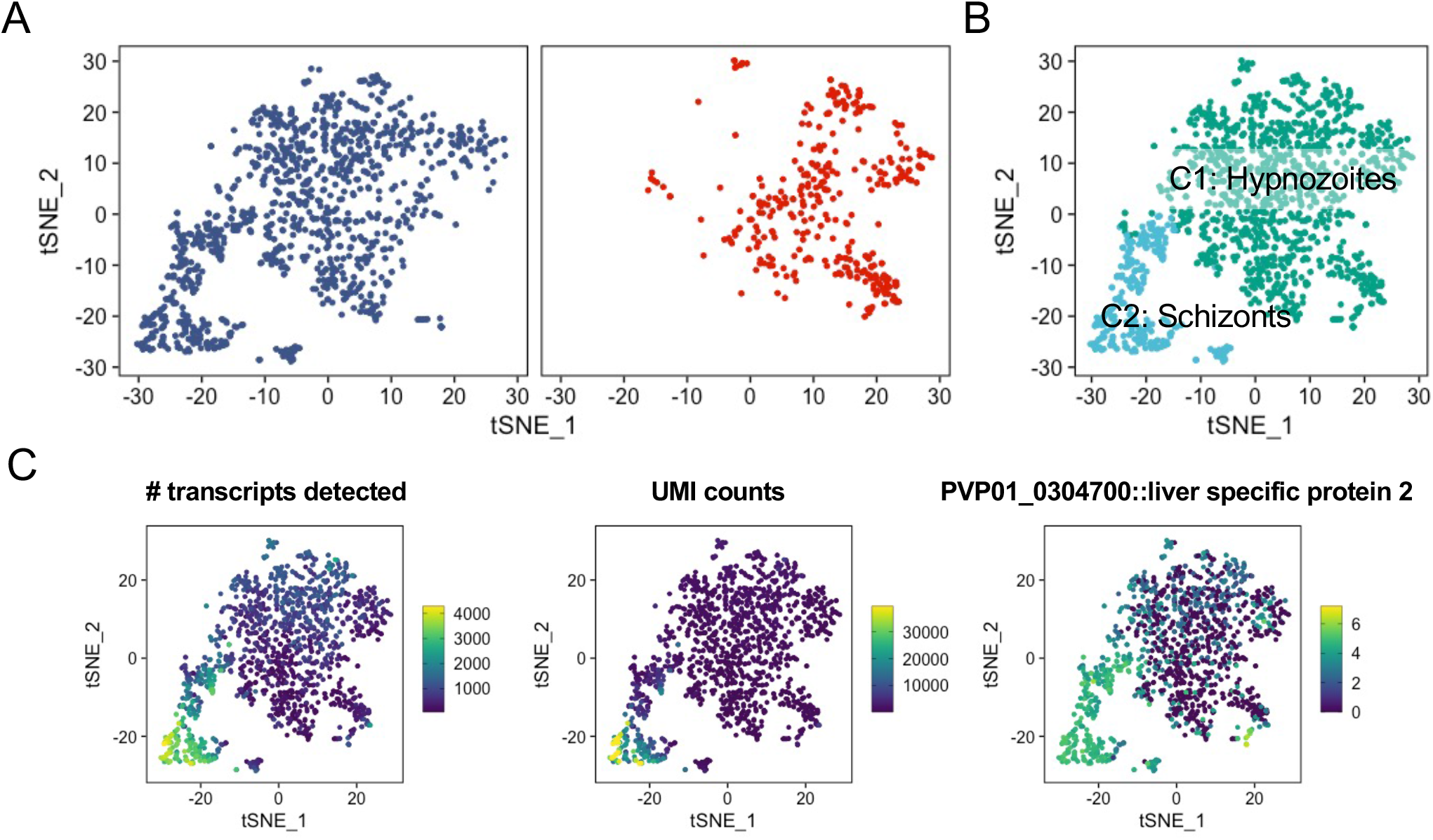
Low dimensional representation of *P. vivax* schizont and hypnozoite transcriptomes. (A) t-SNE plots of integrated *P. vivax* liver stage scRNA-seq data faceted by parasites isolated on day 5 (left) and day 9 (right) post-infection. (B) t-SNE plot of *P. vivax* liver stage data colored by parasites encoding for putative hypnozoites (C1) and schizonts (C2). (C) t-SNE plots of *P. vivax* liver stage scRNA-seq data colored by number of transcripts detected (left), UMI counts (middle), and the expression of the schizont marker, LISP2 (PVP01_0304700) (right). Scales: absolute count (left, middle), normalized expression (right).

Previous scRNA-seq studies in *Plasmodium* spp. has shown gene expression varies with life-cycle stage, with non-replicating forms displaying less expression per cell compared to replicating forms (Howick et al., 2019). The median number of genes detected in schizonts (2323) was greater than the values observed in other single-cell assessments of replicating stages of the parasite’s life cycle (**Figure 2—figure supplement 2**) (Mancio-Silva et al., 2022; Sà et al., 2020). Furthermore, the median number of UMIs per schizont (6905) was also greater in the current dataset versus other reports. The median number of genes detected per hypnozoite (519) and median number of UMIs per hypnozoite (672) were consistent with other single-cell reports assessing non-replicating, uninucleate stages (Bogale et al., 2021; Real et al., 2021; Ruberto et al., 2021a), including recent single-cell assessments of *P. vivax* sporozoites (Ruberto et al., 2021b) and hypnozoites (Mancio-Silva et al., 2022) (**Figure 2—figure supplement 2**). The intact parasitophorous vacuole membrane of hypnozoites in cultures assessed on days 5 and 9 (Figure 1B **and Figure 1—figure supplement 1**), the association between infection rates observed by IFA and the number of transcriptomes assessed, and the per cell metrics typical of *Plasmodium* spp. life stages with low RNA content cells together provided us with strong assurance that the transcriptomes obtained represented viable hypnozoites. These findings show that the transcriptomes of *P. vivax* schizonts and hypnozoites can be distinguished using our experimental workflow.

### Hypnozoites exhibit distinct transcriptional signatures compared to schizonts

We next performed differential gene expression analysis to identify transcriptome-wide differences distinguishing schizonts and hypnozoites. Using stringent inclusion cutoffs (Bonferroni adjusted p value < 0.01 & absolute (average log_2_ fold change [FC]) > 0.5), we found significant differences in the expression of ∼18% (836/4722) of genes detected (**Figure 3—figure supplement 1A; Figure 3— source data 1A**). Relative to hypnozoites, we found an increase in the expression levels of 368 genes in schizonts. As expected, we observed a significant increase of LISP2 gene expression in schizonts (average log_2_ FC: 2.32) compared to hypnozoites. Shortly after invading the hepatocyte, non-relapsing *Plasmodium* spp. liver forms undergo immediate replication. Studies in rodent malaria models have shown that these forms are metabolically active and that various genes involved in energy metabolism are upregulated (Caldelari et al., 2019; Tarun et al., 2008; Toro-Moreno et al., 2020). Consistent with these prior observations, we found an increased transcription associated with glycolysis (PVP01_0816000, PVP01_1229700, PVP01_1244000) and the TCA cycle (PVP01_0106100, PVP01_1332400) in schizonts (**Figure 3—figure supplement 1B; Figure 3—source data 1A**). More broadly, we found increased transcription of various metabolic processes, as well as motility, DNA-, RNA-, and protein-processing (**Figure 3—figure supplement 1C; Figure 3—source data 1A**). Genes encoding for various proteasome subunits displayed greater transcription in schizonts (**Figure 3—figure supplement 2A; Figure 3—source data 1A**). To further show the importance of this protein complex, we tested a known proteosome inhibitor, carfilzomib (Kortuem and Stewart, 2013), in our standard 12-day *P. vivax* small molecule screening assay (Maher et al., 2021), and found it effectively killed schizonts (IC_50_ 288nM) and hypnozoites (IC_50_ 511nM) with some selectivity over host hepatocytes (IC_50_ 3.32 µM) (**Figure 3—figure supplement 2C and D**).

The detection of genes associated with energy metabolism in hypnozoites, albeit at lower levels relative to schizonts, suggested that these forms are also metabolically active (**Figure 3—figure supplement 1B**). Relative to schizonts, we found an increase in expression of 468 genes in hypnozoites (**Figure 3—source data 1A**). As represented in Figure 3A, genes with greater expression in hypnozoites encoded for a diverse set of ontologies—including proteases, membrane proteins, transcription factors, and DNA-/RNA-regulating proteins. Genes encoding for proteases vivapain-1, vivapain-2 (PVP01_1248900, PVP01_0916200), and plasmepsin IV (PVP01_1340900) displayed the greatest differential expression in these forms compared to schizonts (Figure 3A and B**; Figure 3— source data 1A**). While limited research has reported the functionality of these gene products in *P. vivax* (Byoung-Kuk et al., 2004), work in other *Plasmodium* spp. has revealed that they localize to the food vacuole and play an important role in hemoglobin catabolism during the blood stages of the parasite’s life cycle (Dame et al., 2003; Drew et al., 2008).

**Figure 3.**
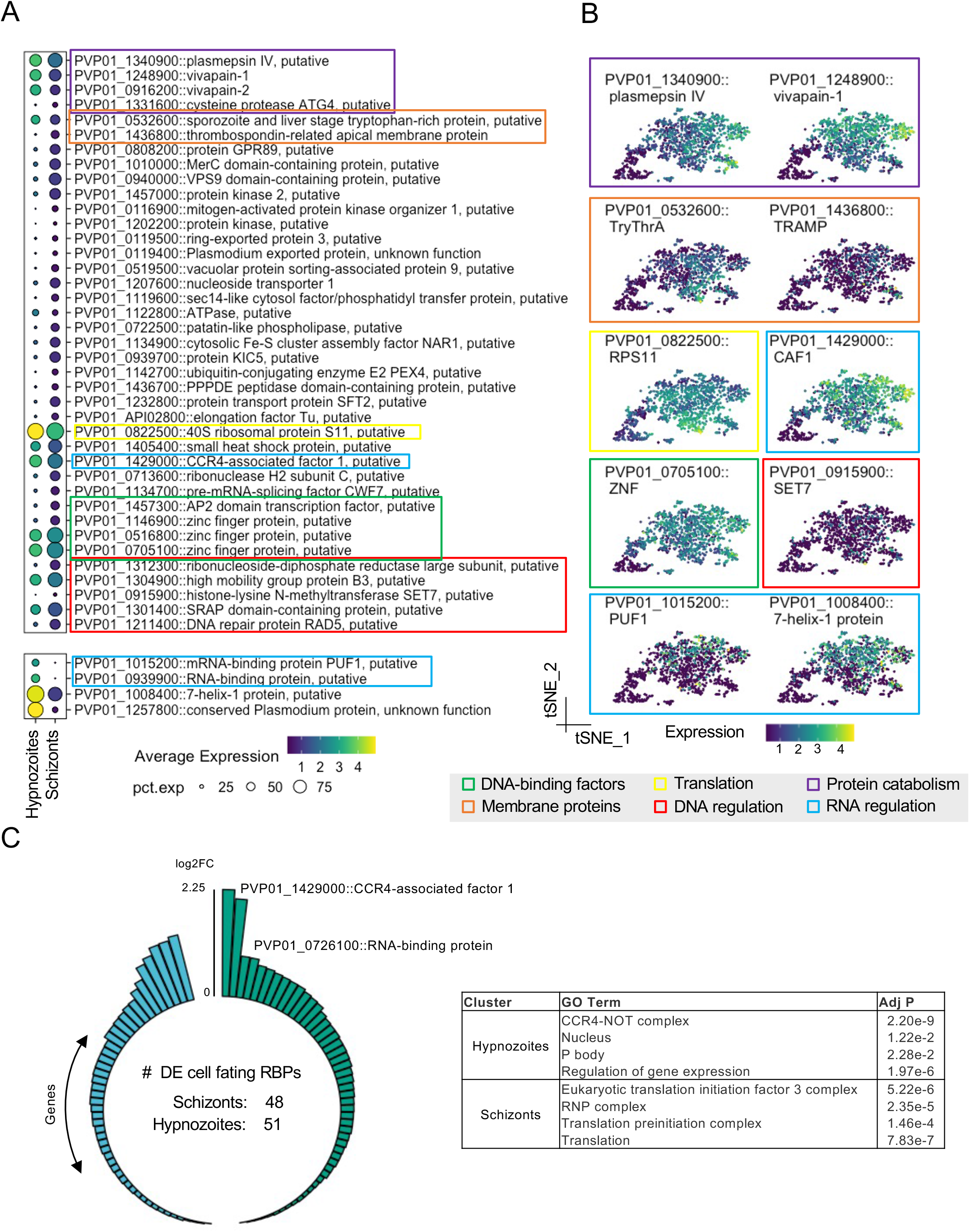
*P. vivax* hypnozoites have distinct transcriptomic signatures. (A) Dot plots showing transcripts that distinguish all hypnozoites from schizonts. The size of the dot corresponds to the percentage of cells that transcript was detected per condition, colored by average expression. Differentially expressed transcripts were identified using Seurat’s FindMarker function (Wilcoxon rank-sum test, Bonferroni adjusted p values < 0.01). (B) tSNE plots of liver stages colored by transcripts displaying greater levels in hypnozoites. Colored boxes in A and B indicate the family or canonical biological process that the transcript is associated with. Scale: normalized expression; pct. exp: percent of cells expressing transcript. (C) Number of transcripts in schizonts and hypnozoites identified as cellular fating RBPs (left) and GO term enrichment of this subset (right). Adj P: Bonferroni adjusted p value.

We also identified genes with greater expression associated with RNA regulatory mechanisms in hypnozoites. Notably, we found greater expression of CCR4 (PVP01_1429000), a gene encoding for a protein with mRNA deadenylase activity and playing a role in mRNA turnover (Tucker et al., 2001) (**Figure 3—source data 1A**). In *Plasmodium* spp., it has been described as important in regulating the expression of genes associated with invasion and egress in blood stages (Balu et al., 2011). RNA-binding proteins (RBPs) also govern gene expression at the post-transcriptional level by transiently storing transcripts until later processing (Hentze et al., 2018). Pumilio domain (PUF) proteins are a major family of RBPs that regulate cellular fating through translational repression in eukaryotes. We found greater expression of the gene encoding for PUF1 (PVP01_1015200) in hypnozoites relative to schizonts (Figure 3A and B; **Figure 3—source data 1A**). Furthermore, we found greater expression of 7-Helix-1 protein (PVP01_1008400) (Figure 3A and B; **Figure 3—source data 1A**)—a stress granule component that interacts with the RNA-binding protein PUF2 and serves to be crucial for protein synthesis in *P. falciparum* (Miao et al., 2010). Numerous studies have shown that *Plasmodium* spp. use various post-transcriptional regulatory mechanisms to coordinate stage transitions and development (Baumgarten et al., 2019; Bunnik et al., 2016; Foth et al., 2011; le Roch et al., 2004; Tarun et al., 2008; Vembar et al., 2016). Our data corroborate these findings and suggest that similar strategies may be used in *P. vivax* to regulate liver stage development. Moreover, these data suggest that RBP-mediated cellular fating and translational repression may be involved in regulating hypnozoite quiescence.

Given the potential role in RBP-mediated cellular fating and translational repression governing hypnozoite quiescence, we performed further analyses focused on RBP-mediated regulation. To this end, we scanned the *P. vivax* genome searching for genes encoding for cellular fating RBPs based on ortholog groups (OGs) of known human cellular fating RBPs (348 RBPs; 227 OGs), identifying 89 OGs (99 RBPs) in *P. vivax.* We detected 48 genes encoding for cellular fating RBPs with greater expression in schizonts (Figure 3C**, left**; **Figure 3—source data 1B**). Notably, we identified eukaryotic translational initiation factors (PVP01_1238100, PVP01_0531200, PVP01_1431600, PVP01_0807600, PVP01_1005600, PVP01_0519700, PVP01_1329200, PVP01_0825000, PVP01_1303500), and splicing factors (PVP01_0716000, PVP01_0521000, PVP01_0804100, PVP01_1111800, PVP01_0607600, PVP01_1012200) suggesting active translational events. Gene Ontology (GO) term analysis revealed that these transcripts were associated with translation initiation (GO:0005852; GO:0033290; GO:0016282) (adjusted p value < 0.05, Figure 3C**, right**; **Figure 3—source data 1C**).

In hypnozoites, we detected 51 genes encoding for cellular fating RBPs with increased expression relative to schizonts (Figure 3C**, left**; **Figure 3—source data 1B**). These genes encode for prominent P body markers, namely DOZI/DDX6 (PVP01_0819400), CCR4-NOT complex proteins (PVP01_1429000, PVP01_1014400, PVP01_1331700, PVP01_1453400, PVP01_0929400), mRNA-decapping enzyme subunit 1 (PVP01_0617200), mRNA-decapping enzyme subunit 2 (PVP01_1409900), UPF1 (PVP01_0805200), UPF2 (PVP01_0724300), UPF3 (PVP01_1226700), PUF2 (PVP01_0526500), PABP (PVP01_1442500), Fibrillarin (PVP01_1341600) and trailer hitch homolog (PVP01_1269800) (**Figure 3—source data 1B**). Genes encoding for cellular fating RBPs in hypnozoite also represented GO terms corresponding to active biological condensate formations, such as the CCR4-NOT complex (GO:0030014; GO:0030015) and the P body (GO:0000932) (adjusted p value < 0.05, Figure 3C**, right; Figure 3—source data 1C**). Together, these findings suggest that P body-mediated cell fating could be important in regulating hypnozoite persistence.

### Transcriptomic heterogeneity exists amongst individual *P. vivax* hypnozoites

Having described the transcriptomic signatures distinguishing schizonts and hypnozoites, we next sought to assess the extent to which gene expression varied amongst hypnozoites. We thus parsed the data into only hypnozoite transcriptomes. In total, 1,147 transcriptomes were assessed (Figure 4A**, left**). Low dimensional representation of the data revealed slight variation between samples collected at days 5 and 9 post-infection (Figure 4A**, right**). After regrouping the data using the same unsupervised clustering algorithm described previously, we identified three clusters (Figure 4B**, right**). The proportion of hypnozoites in each cluster varied across the day of collection: specifically, the proportion of hypnozoites in cluster 3 (C3) was greater in hypnozoites collected at day 9 compared to day 5 post-infection (Figure 4B**, left; Figure 4—figure supplement 1A**). Albeit lower compared to schizonts, we also observed a trend for slightly greater gene expression of LISP2 in C3 relative to the other two clusters (Figure 4C).

**Figure 4.**
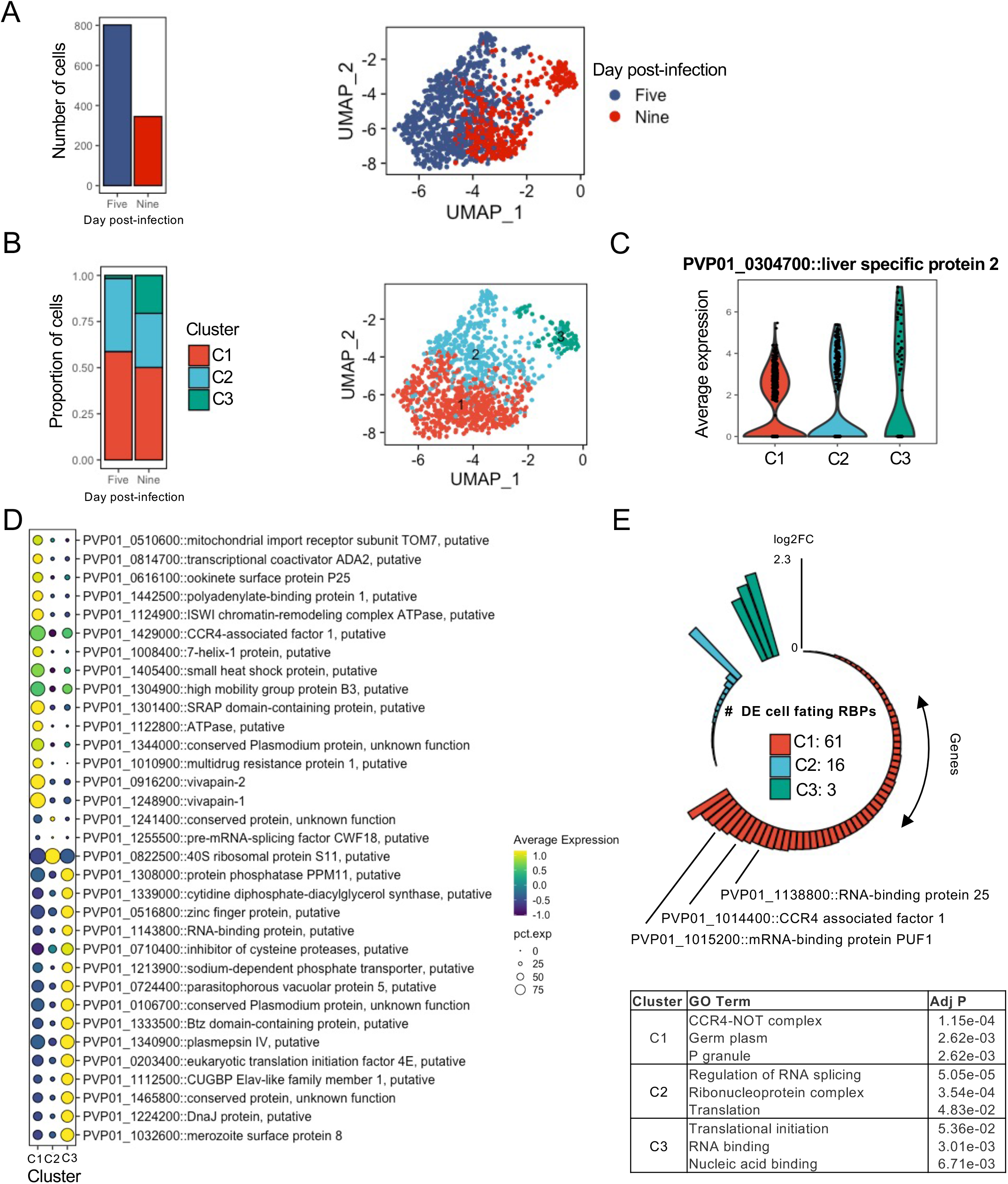
Transcriptomic signatures of activating and persister hypnozoites. (A) Number of hypnozoites assessed (left) and UMAP of cells identified as hypnozoites colored by day of collection. (B) Proportion of cells in clusters on days 5 and 9 post-infection (left) and UMAP of hypnozoites colored by cluster. (C) Violin plot showing the expression of LISP2 in each cluster. (D) Dot plot showing markers that distinguish hypnozoites in each cluster. The size of the dot corresponds the percentage of cells in the expressing the gene of interest, colored by scaled normalized expression of the transcript in each cluster. Marker transcripts were identified using Seurat’s AllFindMarkers function. Wilcoxon rank-sum test, average log_2_FC > 1, Bonferroni adjusted p value < 0.01. (E) Number transcripts in each cluster encoding for cellular fating RBPs sorted by their average log2FC relative to the other clusters (top) and GO Term enrichment of genes in each cluster (bottom). Adj P: Bonferroni adjusted p value.

We next identified markers that define the hypnozoites in each cluster using the FindAllMarkers function in Seurat (Stuart et al., 2019). We defined a marker as a gene displaying greater than 0.5 FC (average log_2_) compared to the other clusters. In total, we detected 477 markers (Bonferroni adjusted p value < 0.05) (**Figure 4—figure supplement 1B**; **Figure 4—source data 1A**). Notable markers in C1 included genes associated with drug resistance (PVP01_1447300, PVP01_1259100, and PVP01_1010900), energy metabolism (PVP01_1435000 and PVP01_1122800), protein modification (PVP01_1248900 and PVP01_0916200), and RNA-binding proteins (PVP01_0939900 and PVP01_0819400); in C2, genes associated with translation (PVP01_1255500 and PVP01_0822500); and in C3, genes associated with translation (PVP01_0213700 and PVP01_0203400), fatty acid metabolism (PVP01_1143000 and PVP01_1022800), protein processing (PVP01_1308000 and PVP01_0927700) and ion transport (PVP01_1407500) (Figure 4D; **Figure 4—source data 1A**).

We next explored the data in the context of cellular fating to elucidate potential patterns in RBP signatures. Among the hypnozoite markers found within our sub-clustering analysis, we identified 80 cellular fating RBPs (Figure 4E**, top; Figure 4—source data 1B**). Approximately 4% (3/80) of the differentially expressed genes in C3 encoded for genes associated with cellular fating. In this cluster, the increased gene expression of eukaryotic translation initiation factor 4E (PVP01_0203400) further suggests active translation events may be occurring. C2 contained 20% (16/80) of the cellular fating RBPs differentially expressed. In this cluster, we found RBPs which are important determinants of cell proliferation from a state of quiescence such as eukaryotic translation initiation factor 5A (PVP01_1303500), polypyrimidine tract binding protein (PVP01_1142400), GTP-binding nuclear protein RAN/TC4 (PVP01_0918300), and transformer-2 protein homolog beta (PVP01_0802200). C1 contained the majority of the differentially expressed genes encoding for RBPs (∼76%, 61/80). Within this cluster, we identified prominent P body markers with cellular fating ability, such as DOZI/DDX6 (PVP01_0819400), CCR4-NOT complex proteins (PVP01_1429000, PVP01_1014400, PVP01_1331700, PVP01_1453400), UPF1 (PVP01_0805200), UPF2 (PVP01_0724300), PABP (PVP01_1442500), Fibrillarin (PVP01_1341600) and trailer hitch homolog (PVP01_1269800). GO analysis revealed enrichment of cellular compartments associated to biological condensate formations, namely CCR4-NOT complex (GO:0030014), P granule (GO:0043186), and germplasm (GO:0060293) (Figure 4E**, bottom; Figure 4—source data 1C**).

These findings, combined with the shift of hypnozoite proportions and increasing level of *LISP2* from C1 to C3, allowed us to infer the phenotype of hypnozoites in these clusters. As these observations likely represent populations of hypnozoites in persisting and activating states, we defined clusters C1, C2, and C3 to represent persisting, early activating, and late activating hypnozoites, respectively. Together, these findings further our understanding of *P. vivax* hypnozoites through characterization of gene usage and putative regulatory mechanisms in activating and persister forms. Furthermore, they highlight a potential role of P body-mediated cellular fating governing persister hypnozoites.

### Strategy used to assess hepatocytes infected with *P. vivax* parasites

After deciphering the identities of *P. vivax* transcriptomes representing schizonts and hypnozoites, we next sought to characterize the transcriptional signatures of hepatocytes infected with either of these forms. To this end, we integrated the data across all the conditions (**Figure 5—supplement 1A and B)** and examined whether distinct sub-populations of infected hepatocytes existed. As highlighted by the low dimensional visualization of the data, we did not find a prominent pattern of infection (Figure 5). Moreover, we detected parasites in hepatocytes with different metabolic states (MacParland et al., 2018), as represented by their transcriptional signatures associated with different hepatocyte zonation programs (**Figure 5—supplement 1C and D**).

**Figure 5.**
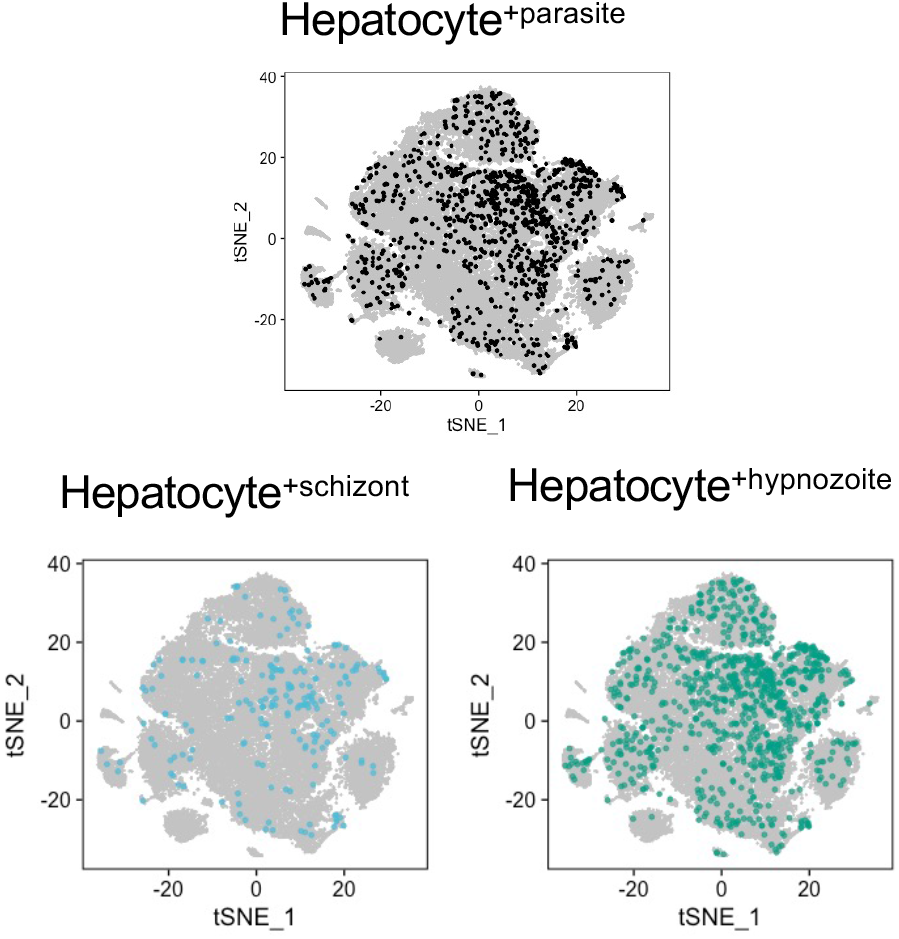
Analysis of host-pathogen transcriptional signatures in *P. vivax-*infected and non-infected hepatocytes. t-SNE plots of hepatocytes faceted by infection status. Grey, noninfected (naive and exposed) hepatocytes; light blue, hepatocytes infected with schizonts; teal, hepatocytes infected with hypnozoites.

### Identification of transcriptional signatures in *P. vivax*-infected hepatocytes

To identify host transcriptional signatures specific to infection status, we performed differential gene expression analysis, comparing infected to non-infected hepatocytes. Among the hepatocytes infected with schizonts, 106 genes displayed greater expression and 16 displayed decreased expression **(Figure 6—source data 1A)**; while in hepatocytes infected with hypnozoites, 365 genes displayed greater expression and 41 genes displayed decreased expression (**Figure 6—source data 1B**) (Bonferroni adjusted p value < 0.05). As depicted in the volcano plots in Figure 6A, the magnitude of change in expression in the infected versus non-infected cells (average log_2_ fold change) was greater in hepatocytes containing schizonts compared to those containing hypnozoites.

**Figure 6.**
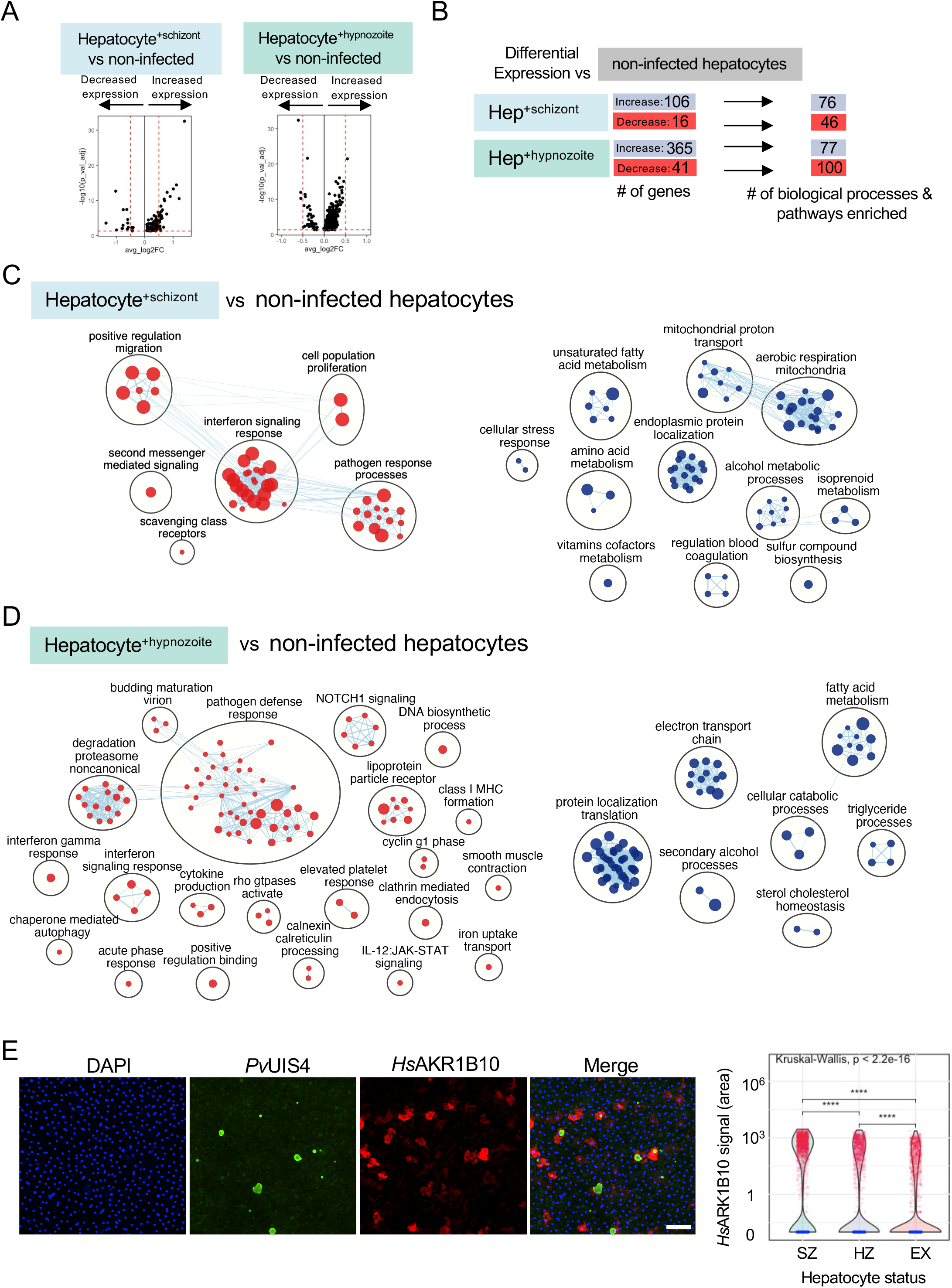
Analysis of host transcriptional signatures in *P. vivax* infected hepatocytes. (A) Volcano plots showing changes in gene expression in infected hepatocytes versus non infected (Wilcoxon rank sum, Bonferroni adjusted p value < 0.05). Positive fold change (FC) values represent genes with greater expression, and negative FC values represent genes with decreased expression. Dashed horizontal red lines: Bonferroni adjusted p value = 0.05; dashed vertical red lines: log_2_FC = 0.5. (B) Summary of differential gene expression data shown in the volcano plots and of the enrichment analyses. Hep: hepatocyte. (C) Enrichment map of cellular processes and pathways associated with hepatocyte infection with *P. vivax* schizonts. (D) Enrichment map of cellular processes and pathways associated with hepatocyte infection with *P. vivax* hypnozoites. For the maps depicted in panels C and D, node size is proportional to the number of genes identified in each gene set (minimum 3 genes/gene set); each node represents a distinct biological process or pathway derived from gene with decreased expression (red) or increased expression (blue) versus non-infected cells; and edges (blue lines) represent the number of genes overlapping between two processes or pathways. (E) Representative confocal image of *P. vivax* parasites on day 12 post-infection of hepatocytes (left). Cells were stained with DAPI (blue), *Pv*UIS4 (green), and *Hs*AKR1B10 (red). White bar represents 100µm. Violin plot displaying the distribution of AKR1B10 signal in hepatocyte cultures infected with *P. vivax* (right). SZ: hepatocyte containing schizont; HZ: hepatocyte containing hypnozoite; EX: hepatocyte not infected but exposed to *P. vivax.* ****: p <= 0.0001.

To gain greater insights into the biological processes and pathways that characterize a schizont-or hypnozoite-infected hepatocyte, we performed enrichment analysis (Raudvere et al., 2019)—scoring 18,594 gene sets against the genes differentially expressed in each of our transcriptional comparisons. When considering both the schizont- and hypnozoite-infected hepatocyte populations, we detected a total of 254 enriched gene sets (Bonferroni adjusted p value < 0.05) (Figure 6B; **Figure 6—source data 1C**). Approximately 18% (45/254) of these gene sets overlapped in hepatocytes infected with either schizonts or hypnozoites (**Figure 6—source data 1D**). These data indicate that while some overlap exists in the gene networks the parasite alters during infection, the majority of processes and pathways altered are specific to schizont versus hypnozoite infected hepatocytes.

Using these gene sets, we then constructed network enrichment maps (Merico et al., 2010) to provide a global overview of the prominent biological themes associated with each infection condition (Figure 6C and D). We detected upregulation of genes associated with pathways related to energy metabolism (oxidative phosphorylation, mitochondrial function, and lipid catabolism) in schizont- and hypnozoite-infected hepatocytes. Analysis of the most differentially expressed genes associated with these pathways included *MT-ND2*, *CHCHD10*, and *AKR1B10* (**Figure 6—source data 1A and B**). Corroborating our transcriptomic data, we found an increase in protein levels of AKR1B10 in infected versus non-infected hepatocytes by IFA (Figure 6E**; Figure 6—figure supplement 1**). AKR1B10 has been implicated in cell survival through its regulation of lipid metabolism and elimination of carbonyls (Martin and Maser, 2009; Qu et al., 2021; Wang et al., 2009) and has primarily been described as a biomarker of cancer (Díez-Dacal et al., 2011; Fukumoto et al., 2005; Jin et al., 2006). More recently, it has been shown that its expression is positively correlated with non-alcoholic fatty liver disease (Govaere et al., 2020). In schizonts, the upregulation of genes linked to fatty acid metabolism is consistent with observations in hepatocytes harboring developing forms from non-relapsing *Plasmodium* spp. (Albuquerque et al., 2009; Itoe et al., 2014). We now reveal that infection with hypnozoites also elicits enrichment of genes associated with fatty acid metabolism (Figure 6D), suggesting that these forms may be scavenging host resources to fuel their persistence.

Previous work assessing *Plasmodium* liver stage biology has revealed that the host is able to detect the parasite (Liehl and Mota, 2012) and that type I interferon plays an integral role in this response (Liehl et al., 2013; Miller et al., 2014). The decrease in expression of genes associated with interferon signaling (REAC:R-HSA-913531, GO:0034340, GO:0034341, GO:0071357, GO:0060337) in hepatocytes infected with either schizonts or hypnozoites suggest that *P. vivax* manipulates the hepatocytes to evade detection (Figure 6C and D**; Figure 6—source data 1C and D**). Furthermore, we found that relative to naïve hepatocytes, uninfected hepatocytes in cultures exposed to *P. vivax* displayed significant downregulation of pathways associated with immune effectors (GO: 0030449, GO:0002697, REAC: R-HSA-166658, R-HSA-168249) (**Figure 6—source data 1E and F**).

In hepatocytes infected with schizonts, we found a decrease in expression of NFKB-regulated genes (*SOD2, TMEM45A, CCL20,* and *TPM1*) (**Figure 6—source data 1A**), consistent with findings that *Plasmodium* spp. release signals to block proinflammatory response (Singh et al., 2007). These findings, in addition to the decrease in the expression of *HLA-B*—a molecule involved in antigen presentation to T cells that can trigger their activation when ‘self’ peptides are not presented—and the neutrophil chemoattractant *CXCL8* suggest that replicating parasites alter hepatocyte signaling pathways to minimize recruitment of innate immune cells and avoid detection by T cells. Similar to hepatocytes infected with schizonts, we found genes associated with interferon signaling downregulated in hepatocytes infected with hypnozoites, namely: *UBB, UBC, HSP90AB1, IFITM3, IFITM2,* and *HLA-A* (**Figure 6—source data 1B**). Furthermore, transcripts associated with the downregulation of class I major histocompatibility complex (*CD81* and *B2M*) were also decreased in these hepatocytes (**Figure 6—source data 1B**). Together, these findings suggest that *P. vivax* infection of hepatocytes may suppress the secretion of various chemokines and proinflammatory cytokines to thwart the recruitment of inflammatory cells to the site of infection while simultaneously downregulating MHC-I to reduce the chances they will be detected if these cells are successfully recruited.

Enrichment analysis linking the changes in gene expression in infected hepatocytes with transcriptional factors revealed distinct sets of putative transcriptional regulators. In total, we detected 35 transcription factors with potential roles in governing the gene expression changes in infected hepatocytes (adjusted p value < 0. 05, Figure 7**; Figure 7—source data 1**). In infected hepatocytes, we found enrichment of transcripts governed by NRF2 (Figure 7**; Figure 7—source data 1)**. The regulation of gene expression through NRF2 in infected hepatocytes may serve to minimize reactive oxygen species in the cell (Hayes and Dinkova-Kostova, 2014). Thus, in addition to the decreased expression of genes associated with inflammatory response (**Figure 6—source data 1C**), NRF2-mediated regulation of antioxidant responsive genes may protect hepatocytes from *Plasmodium*-induced stress and thus serve as a complementary means of minimizing the host inflammatory response.

**Figure 7.**
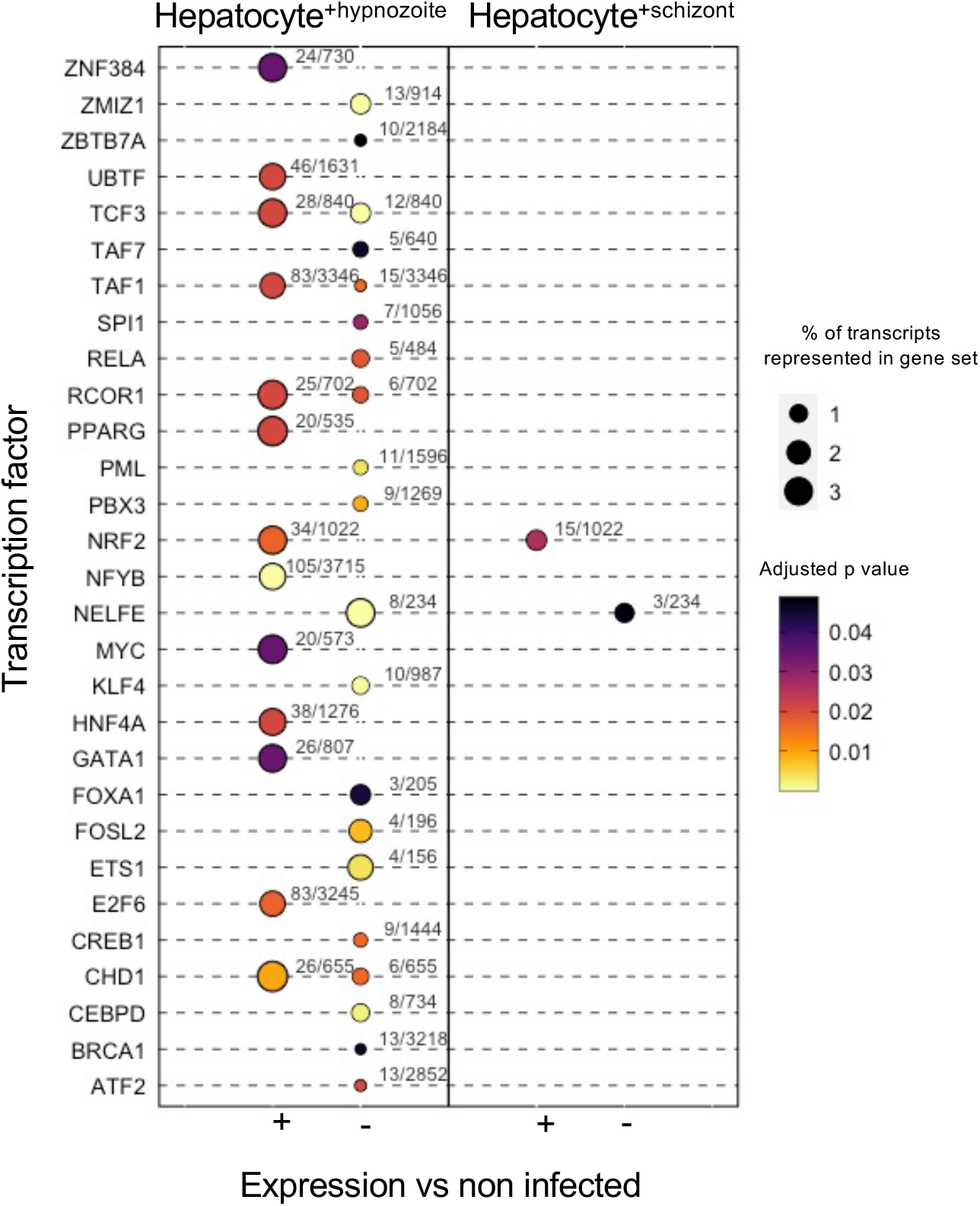
Transcription factor enrichment in hepatocytes infected with schizonts or hypnozoites. Dot plot of transcription factors enriched in Hepatocyte^+hypnozoite^ (left) or Hepatocyte^+schizont^ sets (right). Positive (+) enrichment derived from genes displaying greater expression versus non-infected hepatocytes; negative (-) enrichment derived from genes displaying decreased expression versus noninfected hepatocytes.

## DISCUSSION

The infection of the liver by *P. vivax* constitutes an obligatory step in its life-cycle. *In vivo* studies have shown that within hepatocytes, *P. vivax* can exist in one of two phenotypically-distinct forms: schizonts or hypnozoites (Cogswell, 1992; Mikolajczak et al., 2015). They have also been observed *ex vivo* by infection of hepatocyte monolayers (Gural et al., 2018; Hollingdale et al., 1985; Roth, Maher et al., 2018; Sattabongkot et al., 2006). A key problem hindering the study of *P. vivax* liver stage biology is the difficulty involved in obtaining a sufficient number of infected hepatocytes to make meaningful comparisons between them. We show that by coupling our *in vitro* liver stage model (Maher et al., 2021; Roth, Maher et al., 2018) with a high-throughput single-cell capture technology (Klein et al., 2015; Macosko et al., 2015; Zheng et al., 2017) that transcriptome-wide characterization of *P.* vivax-infected hepatocytes is feasible. The findings provide much needed insight into the molecular signatures of *P. vivax* hypnozoites and the host cells they infect.

Our work corroborates the growing number of *Plasmodium* spp. scRNA-seq data sets, revealing highly variable gene expression patterns across, and within, the various stages of the parasite’s life cycle. Smaller and non-replicating parasites stages (i.e. rings, sporozoites) tend to have fewer genes detected using single-cell approaches. Here, the number of genes detected, as well as the UMIs, per hypnozoite is in line with other single-cell reports assessing the transcriptomic signatures of non-replicating stages of the parasite’s life cycle (Bogale et al., 2021; Howick et al., 2019; Real et al., 2021; Ruberto et al., 2021b), including hypnozoites (Mancio-Silva et al., 2022). Our transcriptome-wide assessment supports the notion that hypnozoites are metabolically active (Adams and Mueller, 2017), demonstrated by the transcription of genes associated with various biological functions including glycolysis and fatty acid metabolism. Furthermore, the expression of genes associated with oxidative stress protection, protein export, mitochondrial respiration, and epigenetic mechanisms such as histone methylation and acetylation, are detectable in hypnozoites, suggesting these parasites are both viable and actively transcribing genes to survive.

We also found that hypnozoites exist in various transcriptomic states. We speculate that these differences might represent a spectrum of phenotypes, from persisting to activating. Interestingly, no loss in gene diversity or expression was found in hypnozoites associated with cluster C1 (persisters) in our hypnozoite-directed analysis, suggesting this subpopulation encodes for viable, distinct forms. In the persister hypnozoite cluster, the differential expression of genes encoding for cellular fating RBPs supports the hypothesis that these forms may use post-transcriptional mechanisms to control gene usage (Mueller et al., 2019). The biological significance of the enrichment of genes associated with translational repression, found exclusively in a subpopulation of hypnozoites (C1), suggests this mechanism may govern quiescence, and warrants further investigation.

Discerning between non-replicating, uninucleate forms and dead parasites poses a challenge. The absence of a validated hypnozoite viability marker further complicates assessments. These limitations notwithstanding, the data reported here seem to suggest that the transcriptomes assessed represent viable parasites. Similar to a recent study using a non-relapsing rodent malaria model (Afriat et al., 2021), we inferred parasite viability based on visual assessment of fixed cell cultures using an immunofluorescence assay (IFA). For each scRNA-seq library processed, we dedicated cultures from the same infection to provide a visual snapshot of the parasite viability at each end point. While Afriat and colleagues (2021) define abortive forms as displaying “vacuole breakdown,” upon inspection of our cultures, we did not find indications of abortive or dead parasites. Moreover, the parasites morphologically identified as hypnozoites were highly similar in morphology to those described in humanized mice (Mikolajczak et al., 2015) and *in vitro* (Roth, Maher et al., 2018). More specifically, they displayed positive *Pv*UIS4 staining for an intact parasitophorous vacuole membrane and a distinctive prominence. These forms have been shown to activate *in vivo* (Mikolajczak et al., 2015) as well as *in vitro* (Maher et al., 2020). Additionally, they have been shown to be susceptible to treatment with primaquine in combination with chloroquine and ionophores, implying they were alive to begin with (Maher et al., 2021). At the molecular level, we did not find consistencies between our data and the transcriptomic signatures of “abortive” *P. berghei* liver forms reported by Afriat and colleagues (2021). For example, orthologous expression of genes encoding for HSP90, HSP70, and HOP—three markers defining abortive parasites in their dataset—displayed greater expression in the schizont cluster relative to hypnozoites. These observations further support the hypothesis that hypnozoite transcriptomes assessesed in this study were derived from viable parasites.

We highlight distinct transcriptomic signatures of infected versus non-infected hepatocytes, and show that signaling pathways associated with energy metabolism, antioxidant stress, and immune response are altered in infected hepatocytes. The extent to which individual hepatocytes respond to *P. vivax* schizonts and hypnozoites has previously been unknown. A bulk RNA sequencing approach has been used to identify pathways altered in hepatocytes in response to *Plasmodium* infection (LaMonte et al., 2019); however, it is limited insofar as it averages gene expression from millions of cells, obscuring any parasite-specific effects on an infected cell. Targeted approaches have revealed *P. vivax* liver forms recruit host factors to facilitate growth (LaMonte et al., 2019; Posfai et al., 2020), but how certain infected hepatocyte populations contribute to disease transmission, and how hepatocyte subpopulations support development, are intriguing topics for future studies (Vijayan et al., 2021). Our data identify potential host factors essential for *P. vivax*’s successful liver stage development. Whether certain hepatocyte phenotypes influence the development, especially activation of hypnozoites into schizonts, requires further investigation.

Several observations suggest protein degradation may be important for the parasite’s liver stages. Targeting mechanisms related to this process may therefore be an effective means of killing these forms. For instance, the ability of hepatocytes to uptake and break down hemoglobin (Bissell et al., 1972) supports that this protein substrate could be an essential amino acid source for the parasite. Our data suggest that several mechanisms may allow *P. vivax* to utilize the amino acids derived from this protein; or more broadly, any protein substrate. In hypnozoites, the increased expression of genes encoding for proteases, such as vivapains, supports the idea that these parasites digest host cytosolic proteins. The role of these proteins as hemoglobinases have been defined extensively in *P. falciparum* blood stages (Drew et al., 2008; Rosenthal, 2011; Salas et al., 1995). A similar role has been described for *P. vivax* (Byoung-Kuk et al., 2004), but there is limited information about their significance in the liver stages. Interestingly, a recent report using a complementary single-cell capture technology (Hughes et al., 2020) also found that these genes display greater expression in hypnozoites relative to schizonts (Mancio-Silva et al., 2022). Vivapains are highly homologous to cathepsin protease L (TgCpl) in the related apicomplexan *Toxoplasma gondii.* In these parasites, TgCpl is localized to its plant-like vacuolar compartment—a lysosome-like organelle containing a variety of proteases (Parussini et al., 2010)—and one of its functions is to aid in the digestion of host-derived proteins (Dou et al., 2014). Genetic or chemical disruption of TgCpl negatively impacts the viability of bradyzoites, a persisting form of this parasite (di Cristina et al., 2017). Given these orthologous proteins’ roles in *P. falciparum* and *T. gondii*, we speculate that the expression of vivapains may support the persistence of *P. vivax* hypnozoites.

Persistence of hypnozoites is contingent on the longevity of the hepatocyte in which it resides. The increased expression of genes associated with antioxidant stress (Scarpulla, 2002) is consistent with the hypothesis that hypnozoite infection induces changes in the hepatocyte to promote cell survival, in which NRF2 could be playing a regulatory role (Bindschedler et al., 2022). One host gene upregulated in infected hepatocytes validated by IFA, *AKR1B10*, is regulated by NRF2, which activates protective pathways in response to oxidative stress (Tebay et al., 2015). We also detected genes displaying greater expression in infected hepatocytes regulated by NFYB, a transcription factor important for cell proliferation, mitochondrial integrity, and cellular longevity (Tharyan et al., 2020). At day 5 post-infection, the increased expression of mitochondria-related genes in hepatocytes infected with schizonts may be required to meet the parasite’s energy demands during replication, but the enrichment of mitochondria-related genes in cells infected with hypnozoites could also aid in ensuring longevity of the hepatocyte. In addition to these positive regulators of cell survival, we found a decreased expression of genes associated with cell death such as *UBB, UBC,* and *HSP90AB1*, potentially regulated by CEBPD, a transcription factor with a role in cell death (Tsai et al., 2017; Wang et al., 2019).

Persistence of hypnozoites also requires evading host detection. Our analyses indicate the parasite alters the host hepatocyte to evade detection by immune cells. First, there is a decrease in expression of genes encoding for chemoattractants (*CCL20, CXCL8, IL32*) in infected hepatocytes, and an increase in expression of PTGR1, a negative regulator of the chemotactic factor leukotriene B4 (Yokomizo et al., 1993). These changes would result in decreased secretion of chemokines that recruit innate immune cells to the liver, thereby reducing recruitment of adaptive immune cells like CD8 T-cells that kill infected hepatocytes (Chakravarty et al., 2007). Interestingly, there was also a decrease in expression of MHC-1 molecules (*HLA-A and -B*). The presentation of parasite peptides to be detected by surveying CD8 T-cells would therefore be reduced (Neefjes et al., 2011), creating an additional layer of protection should innate immune cells be recruited, for example upon lysis of the hepatocyte by schizonts. We propose that *P. vivax* infected hepatocytes are re-programmed to prevent recruitment of innate immune cells on the one hand, and to reduce the likelihood of their detection by the adaptive immune system on the other, all to enable progress through schizogony and for hypnozoites to persist undetected. Understanding the mechanisms leading up to this warrant further inversitgation, as their reversal may lead to improved clearance of hypnozoites, thus reducing relapses. Indeed, a vaccine would need to decrease relapses to have an impact on *P. vivax* transmission and disease (White et al., 2017), but the decrease of MHC-I expression in infected hepatocytes may mean that standard focuses to generate CD8 and CD4 polyfunctional T-cells in the liver may not effectively eliminate *P. vivax*. Recent attempts to test anti-relapse vaccine candidates with *P. cynomolgi* did not prevent relapses despite higher inducing IFNγ producing cells, a common metric used as a correlate of protection (Kim et al., 2020), indicating that alternative strategies may be needed.

Our transcriptome-wide characterization of schizonts, hypnozoites, and infected hepatocytes highlights transcripts altered during infection, and provides a framework for examining the role of infection dynamics of *P. vivax* liver stages. By examining two early liver stage time points at single-cell resolution, our study lays out a proof of principle and a technical advance demonstrating the importance of *in vitro* models to shed light on the molecular basis of *P. vivax* liver stage development and infection. To conclude, the overall aim of our research was to elucidate the molecular signatures of *P. vivax* liver infection, a better understanding of which will aid in facilitating the development of new therapeutics to ultimately reduce transmission. We anticipate that interventions could involve parasite- and/or host-directed small molecule therapies, both of which our group is actively pursuing. The comprehensive mapping of the transcriptional landscape of both the host and *P. vivax* described here provides an important framework for further investigation.

## MATERIALS AND METHODS

### Ethics statement

All research procedures were reviewed and approved by the Cambodian National Ethics Committee for Health Research (approval numbers: #113NECHR, #104NECHR and #089NECHR).

### Blood samples, mosquitos, and infections

The complete protocol used for mosquito rearing an infection has been published (Maher et al., 2021). In brief, blood samples from symptomatic patients infected with *P. vivax* were collected at local health facilities in Mondulkiri province in Cambodia from March 2020 to October 2021. Following a *P. vivax* gametocyte-containing blood meal, *An. dirus* mosquitoes were maintained at ambient temperature on a 12:12 light: dark cycle and fed 10% sucrose + 0.5% PABA solution. *An. dirus* found positive for *P. vivax* oocysts at six-days post-feeding were transported to the Institut Pasteur Cambodia Insectary Facility in Phnom Penh, Cambodia where they were maintained under the same conditions described above.

### Sporozoite isolation and primary human hepatocyte infections

The complete protocol used for seeding primary human hepatocytes, harvesting sporozoites, and infecting cultures has been published (Maher et al., 2021). In brief, *P. vivax* sporozoites were isolated from the salivary glands of female *An. dirus* mosquitoes 18 days after an infectious blood-meal. Primary human hepatocytes (Lot BGW, BioIVT) were seeded 2 days before infection with 30,000 sporozoites (replicate 1), or 15,200 sporozoites (replicate 2) per well. Media was replaced with fresh CP media (BioIVT) containing antibiotics (penicillin, streptomycin, and gentamycin) the day after infection and every 2-3 days thereafter. Hepatocytes were treated with 1 µM MMV390048 on days 5, 6, and 7 post-infection to kill replicating schizonts, resulting in cultures with solely hypnozoites at 9 days post-infection. At days 5 and 9 post-infection, cultures were processed for single-cell partitioning (described in the following section) or immunofluorescence assay (IFA). For the latter, hepatocytes were fixed with 4% paraformaldehyde in 1X PBS. Fixed cultures were then stained overnight at 4°C with recombinant mouse anti-*P. vivax* UIS4 antibody (Schäfer et al., 2018) diluted 1:25,000 in a stain buffer (0.03% Triton X-100) and 1% (w/v) BSA in 1X PBS. The following day, cultures were washed three times with 1X PBS and then stained overnight at 4°C with rabbit anti-mouse AlexaFluor ™488-conjugated antibody (Thermo Fisher) diluted 1:1,000 in stain buffer. Cultures were washed three times with 1X PBS and counterstained with 1 mg/mL Hoechst 33342 (Thermo Fisher) to detect parasite and host DNA. High content imaging was performed on a Lionheart imaging system (Biotek). Quantification of the number of nuclei and parasite liver forms was performed using Gen5 high content analysis software (Biotek). Confocal images were obtained using an ImageXpress Imaging System (Molecular Devices).

### Single-cell partitioning, library preparation, and sequencing

Hepatocyte cultures infected with *P. vivax* were processed prior to- and post-MMV390048 treatment (5- and 9-days post-infection, respectively). Two replicate experiments, defined as an independent plating of hepatocytes infected with sporozoites from a unique field isolate, were performed several months apart. Additionally, and as a negative control, one replicate of hepatocyte cultures never exposed to *P. vivax* (naïve) were processed similarly to those that were infected. At each endpoint for each replicate, approximately 640,000 primary human hepatocytes (16 wells of a 384-well plate each having approximately 10,000 cells) were treated with trypsin (Corning 25-053-Cl) to facilitate their detachment from the plate. Once detached, trypsin was inactivated with 1:1 complete media and the cell suspension was collected into a 5mL protein-low bind centrifuge tube (Eppendorf). To remove residual trypsin, hepatocytes were washed via three sequential rounds of pelleting (50 x g, 3 min, 4°C) and resuspending in 1X PBS containing 0.1% BSA. Cells were then passed through a 35µm strainer (Falcon 352235) to minimize or remove hepatocyte clusters. Prior to partitioning cells on the Chromium controller (10x Genomics), hepatocytes were counted and the cell concentration was assessed using a hemocytometer. Approximately 10,000 hepatocytes were loaded on a Chromium Chip B (10x Genomics) as per manufacturer’s recommendations. Chips containing hepatocyte suspensions, Gel Beads in Emulsion (GEMs), and reverse transcription reagents were loaded into the Chromium controller (10x Genomics) for single-cell partitioning and cell barcoding. Barcoded cDNA libraries were generated according to the Chromium Single Cell 3’ gene expression protocol (version 3). In total, 6 cDNA libraries were generated (two replicates at 5 days post-infection, two replicates at 9 days post-infection, one uninfected control for day 5 cultures, and one uninfected control for day 9 cultures). Libraries were loaded on individual flow cell lanes and sequenced at a depth of 400 million reads using a HiSeq X Ten platform (Illumina) at Macrogen (Seoul, Korea).

### Computational analysis of single-cell RNA sequencing data

Quality control checks on the raw sequencing data were performed using FASTQC (Andrews, 2010). In a manner similar to other host-pathogen single-cell studies (Ren et al., 2021; Szabo et al., 2021), a multi-species transcriptome index containing the transcriptomes for *P. vivax* P01 (PlasmoDB.org, version 51) and *H. sapiens* GRCh8 (Ensembl, version 103) was created using the ‘ref’ function in kb-python (0.26.3)— a python package that wraps the kallisto | bustools single-cell RNA-seq workflow (Bray et al., 2016; Melsted et al., 2021). The *P. vivax* P01 transcriptome file used for the generation of the multi-species index contained gene regions encoding for UTRs (Siegel, Chappell et al., 2020). Next, using the same package, reads were pseudoaligned and counted using the ‘count’ function with parameters suitable for sequencing data generated from version 3 of 10x Genomics’ 3’ gene expression technology: kb count -i human_Pv_index.idx -g t2g.txt -x 10xv3 -t8 -o . /path/to/read/10xV3sequencesread1.fastq.gz path/to/10xV3sequencesread2.fastq.gz. After performing kb count on the sequencing outputs from the 6 single-cell libraries, the resulting unfiltered cell-gene count matrices were read into R (v 4.1.1) for further host and parasite-specific processing described in the following sections.

### Filtering and normalization of scRNA-seq count matrices

To distinguish between droplets containing cells and those only with ambient RNA, we used DropUtils’ emptyDrops function (Lun et al., 2019) with the lower total UMI count threshold (at or below which all barcodes are assumed to correspond to empty droplets) set to 1,000. The droplets with cells (FDR < 0.001), more specifically, those containing barcoded transcripts derived from either human, parasite, or a combination of the two—were kept for downstream processing.

### Assessment of *P. vivax* liver stage transcriptomes

For the analyses of *P. vivax*, we filtered out the transcripts derived from human, leaving only transcripts derived from the parasite to be assessed. All samples were processed independently. For each sample, cells with less than 60 genes detected were removed. Next, we filtered out genes with low counts, retaining genes with 10 or greater UMIs across all cells. Post cell and gene filtering, the data were normalized using Seurat (Stuart et al., 2019) ‘LogNormalize’ function with the default parameters selected.

#### Data integration and clustering

The data were integrated using Seurat’s SCTransform function (Hafemeister and Satija, 2019) with the following parameters indicated: variable.features.n = NULL, variable.features.rv.th = 1.3. Clustering was performed using the Seurat functions FindNeighbours (dims = 1:20) followed by FindClusters (resolution = 0.1, algorithm = 4 for all data or resolution = 0.3, algorithm = 4 for hypnozoite filtered data).

#### Differential gene expression analysis

To detect differentially expressed genes, the Seurat function FinalAllMarkers and FindMarkers were used. For differential gene expression testing using the FindMarkers function, we used the following parameters: logfc.threshold = 0.0, ident.1 = ‘1’, ident.2 = ‘2’, test.use = “wilcox”, min.pct = 0, only.pos = F, assay = “RNA”. For differential gene expression testing using the FindAllMarkers function, we used the following parameters: test.use = “wilcox”, only.pos = T, assay = “RNA”, log2FC = 0.0. For the schizont versus hypnozoite comparisons, genes with a Bonferroni adjusted p value < 0.01 & absolute average log2 FC) > 0.5 were used RBP and GO analyses. For the hypnozoite versus hypnozoite comparisions, genes with a Bonferroni adjusted p value of < 0.05 and absolute average log2 FC > 0.5 were used for RBP and GO analsysis.

#### RBP analyses

RBPs with cell fating potential reported in humans (Decker and Parker, n.d.; Gerstberger et al., 2014; Guallar and Wang, 2014; Voronina et al., 2011; Youn et al., 2019) were manually curated. The orthologues of these cell fating RBPs in *P. vivax* were obtained using OrthoMCL (Release-6.8)(Chen et al., 2006). Within the differentially expressed genes, genes encoding for cell fating RBPs were filtered for further assessment.

#### GO enrichment analyses

GO enrichment analysis was performed on the PlasmoDB (v55) website with the following input parameters selected: P value cutoff = 0.05; Limit to GO Slim terms = No; Evidence: Computed, curated. Enrichment from GO cellular compartment (CC), biological processes (BP), and molecular function (MF) were all assessed. Gene with an Bonferroni adjusted p value < 0.05 were considered significantly enriched.

### Assessment of human hepatocyte transcriptomes

For the analyses of hepatocytes, we removed all transcripts derived from *P. vivax*, leaving only the transcripts derived from hepatocytes. We retained transcripts detected in at least 30 cells. Next, cells with less than 1,500 UMIs were removed from further analyses. Post-cell and gene filtering, the data from each replicate were normalized using Seurat’s ‘LogNormalize’ function with the default parameters.

#### Data integration and low dimensional reduction

Filtered, normalized data matrices were integrated in a manner described in in the Seurat (version 4) vignette, *Introduction to scRNA-seq integration*, available on the Sajita Lab’s website (https://satijalab.org/seurat/articles/integration_introduction.html). Briefly, integration features were identified in each replicate using the ‘SelectIntegrationFeatures’ function with the nFeatures parameter set to 3000. Next, we used ‘PrepSCTIntegration’ to prepare object list containing the 3000 features for integration. We then used ‘FindIntegrationAnchors’ to identify a set of anchors between datasets with the following parameters specified: normalization.method = “SCT”, anchor.features = “features”, reference = naïve hepatocytes day 5). Last, using these anchors, the six datasets were integrated using the ‘IntegrateData’ function with the following parameters specified: normalization.method = “SCT”. After integrating the data, we perform dimensionilty reduction using the ‘RunPCA’ function, followed by ‘RunTSNE’ with 1:30 dimensions selected.

#### Gene Ontology and Reactome enrichment analyses

Enriched biological processes and pathways were determined using gProfiler (Raudvere et al., 2019) with the following options selected: ordered query, statistical domain scope—only annotated genes, significance threshold—g:SCS threshold, user threshold—0.05, Gene Ontology—GO biological processes, No electronic GO annotations, biological pathways—Reactome. Transcripts that displayed significant changes in expression in infected hepatocytes versus non-infected hepatocytes were used as input. EnrichmentMap (v 1.1.0) (Merico et al., 2010) for CytoScape (v 3.8.2) (Shannon et al., 2003) was used to create the enrichment networks. Significantly enriched GO biological processes and Reactome pathways (adjusted p value < 0.05) were used as input. Pathways were clustered and annotated using the AutoAnnotate function (Kucera et al., 2016). Node size corresponds to the number of transcripts in the corresponding set, and edge widths correspond to the number of transcripts shared between sets.

#### Transcription factor enrichment analysis

We downloaded curated sets of known transcription factors from the website (http://amp.pharm.mssm.edu/Enrichr/) (Kuleshov et al., 2016). Transcription factor—target gene sets were obtained by ChIP-X experiments from the ChEA (Lachmann et al., 2010) and ENCODE (Feingold and Pachter, 2004) projects. TF-target relationships using Chip-seq data from ENCODE and CHEA were used to identify overrepresented pathways and TFs. Differentially expressed genes (Bonferroni adjusted p value < 0.05) in each condition versus non-infected cells were used as inputs. Gene sets with an q value < 0.05 were considered significantly enriched.

### Carfilzomib dose-response assessment

A carfilzomib dose-response against *P. vivax* liver forms was obtained from three independent experiments, in which a different *P. vivax* clinical isolate was allowed to infect BGW human hepatocytes. The 12-day assay was used (Maher et al., 2021). All independent experiments resulted in sufficient hypnozoite formation within each well for dose-response calculations. Liver stage parasite growth metrics and compound dilutions were loaded into CDD Vault (Collaborative Drug Discovery) and growth data normalized such that zero (negative) inhibition was set as the average of DMSO wells and 100% (positive) inhibition was set to the effective doses of a nigericin control. Normalized values were then combined in Graphpad Prism to fit a dose-response curve and calculated IC_50_ values from all replicates.

### Quantification of AKR1B10 protein expression

To analyze AKR1B10 expression and localization, *P. vivax*-infected primary human hepatocytes were fixed 12 days post-infection. Five independent hepatocyte infections were used. Infected wells were then permeabilized for 20 minutes with 0.2% TritonX-100, washed thrice with 1X PBS, and blocked in 3% BSA for one hour. Primary antibodies rabbit anti-HsAKR1B10 (PA5-30773, Thermo Fisher) and mouse anti-*Pv*UIS4 were sequentially added, diluted 1:100 and 1:500, respectively. Antibodies were incubated overnight at 4°C and washed 3 times with 1X PBS before addition of 1:400 Alexa-Fluor 568 donkey anti-rabbit for *Hs*AKR1B10 or 1:400 AlexaFluor 488 goat anti-mouse for *Pv*UIS4. Secondary antibodies were incubated overnight at 4°C. Cells were washed thrice with 1X PBS before adding 0.5 mg/mL DAPI for 10 minutes. Cells were washed 3 final times with 1X PBS before image collection with an ImageXpress inverted confocal microscope (Molecular Devices). Image analysis was completed using MetaXpress analysis software (Molecular Devices).

### Contact for reagent and resource sharing

Further information and requests for resources and reagents should be directed to and will be fulfilled by the Lead Contact, Dennis Kyle.

### Data availability

All raw sequencing data have been deposited in the European Nucleotide Archive at European Molecular Biology Laboratory European Bioinformatics Institute (www.ebi.ac.uk/ena/) under accession number PRJEB52293. Scripts containing the code used to process the single-cell RNA seq data are available on GitHub at: https://github.com/AnthonyRuberto/Pv_LS_singleCell. Archived scripts (Shell and R) and output files as at time of publication will be made available on Zenodo.

## Acknowledgements

We thank the patients of Mondulkiri Province, Cambodia, for participating in this study. We thank the Institut Pasteur du Cambodge’s field site manager (Saorin Kim) for logistical assistance, insectary staff (Makara Pring, Koeun Kaing, Nora Sambath) for the *An. dirus* mosquito colony maintenance, and laboratory staff (Eakpor Piv, Chansophea Chhin, Sreyvouch Phen, Chansovandan Chhun, Sivcheng Phal, Baura Tat) for assistance with the mosquito dissections and the *in vitro* assays.

High-content imaging data was produced in part by the Biomedical Microscopy Core at the University of Georgia, supported by the Georgia Research Alliance. A.R.J., B.B. and I.M. acknowledge support from the Victorian State Government Operational Infrastructure Support and Australian Government National Health and Medical Research Council Independent Research Institute Infrastructure Support Scheme. This work was also supported by the Agence Nationale de la Recherche (ANR-17-CE13-0025 to A.A.R and I.M.); an Australian National Health and Medical Research Council Leadership Fellowship (APP1194330 to A.R.J.); the Bill & Melinda Gates Foundation (OPP1023601 to D.E.K.); and Medicines for Malaria Venture (RD/2017/0042 to B.W. and A.V., RD/16/1082 and RD/15/0022 to S.P.M. and D.E.K.).

A.A.R., S.P.M., A.V., I.M., B.W. and D.E.K conceived and designed the study; A.A.R, S.P.M. and A.V. performed the experiments; A.A.R., C.B., B.B. and C.J. curated the data; A.A.R., S.P.M., C.B., B.B., A.R.J. and C.J. performed the formal analyses; S.P.M., A.V. and I.M., B.W. and D.E.K. coordinated the research activities; S.P.M., A.V., I.M., B.W. and D.E.K. provided resources; S.P.M. and D.E.K. acquired funding for the work; A.A.R wrote the initial draft of the manuscript; all authors reviewed and made contributions to the final version of the manuscript.

---

**Figure 1—figure supplement 1**

**Assessment of *P. vivax* liver stages using high-content imaging.** Representative high content images of primary hepatocytes infected with *P. vivax* on day 5 (left) and day 9 (right) post-infection. Images were obtained from one well from a 384-well plate with a 4x objective. Inset: one field of view (orange box) from the same well captured with a 20x objective. Cells were stained with DAPI (blue) and *Pv*UIS4 (green). White arrow: liver form assigned as a hypnozoite, yellow arrow: liver form assigned as a schizont. White bar represents 1mm, grey bar represents 200µm. Images are representative of the second biological replicate used in the study.

**Figure 2—figure supplement 1**

**Metrics associated with *P. vivax* liver stage scRNA-seq data analyses.** (A) Number of *P. vivax* liver stage transcriptomes assessed post-cell and gene filtering. (B) Scatter plots displaying the number of genes detected versus total number of UMIs for each parasite transcriptome assessed. Red dashed vertical line (60) represents the cut-off used to filter viable from problematic (dead/dying/poorly captured) cells. (C) Number of *P. vivax* liver stage parasites assigned to cluster 1 (hypnozoites) and cluster 2 (schizonts). SZ: schizont; HZ: hypnozoite; dpi: day post-infection.

**Figure 2—figure supplement 2**

**Comparision of per cell metrics with other *P. vivax* scRNA-seq data.** Violin plots showing the distribution of genes detected in the current dataset, and other single-cell gene expression assessements of *P. vivax* liver stages (Mancio-Silva et al., 2022), sporozoites (Ruberto et al., 2021b), and blood-stage parasites (Sà et al., 2020). Replicates in sporozoite and blood-stage datasets represent unique 10x Genomics’ scRNA-seq library preparations. Blood-stage scRNA-seq metrics were obtained after realignment of sequencing data to an updated *P. vivax* transcriptome including UTRs as performed previously (Ruberto et al., 2021b). N = number of single-cell transcriptomes assessed for each sample. Light blue dashed horizontal line: median number genes detected or UMIs in schizonts obtained in the current dataset; teal dashed horizontal line: median number genes detected or UMIs in hypnozoites obtained in the current dataset. Day denotes day post-infection.

**Figure 2—source data 1**

**Summary statistics and cell and gene filtering metrics for each single-cell RNA library.**

**Figure 3—figure supplement 1**

***P. vivax* schizonts and hypnozoites have distinct transcriptomic signatures.** (A) Scatterplot showing average log_2_ fold change versus the difference in the proportion of cells the transcript is detected in clusters encoding for hypnozoites and schizonts. Negative values: greater detection in hypnozoites; positive values: greater detection in schizonts. (B) Violin plots displaying the expression of genes encoding for TCA- and glycolysis-related proteins. (C) Dot plots showing transcripts with decreased expression in hypnozoites relative to schizonts. The size of the dot corresponds to the percentage of cells expressing the gene, colored by average expression. Differentially expressed transcripts were identified using Seurat’s FindMarkers function. Wilcoxon rank-sum test, Bonferroni adjusted p values < 0.01. Scale: normalized expression; pct. exp: percent of cells expressing the gene. ldh: L-lactate dehydrogenase; gapdh: glyceraldehyde 3-phosphate dehydrogenase; atp-synthC: ATP synthase subunit C; gdh1: NADP-specific glutamate dehydrogenase 1.

**Figure 3—figure supplement 2**

**Carfilzomib inhibits *P. vivax* schizonts and hypnozoites.** (A) Violin plots displaying the distribution of expression of select genes encoding for proteasome subunits in schizonts and hypnozoites. (B) Table of proteasome-associated transcripts differentially expressed between schizonts and hypnozoites. Positive average log2FC: higher expression in schizonts. (C) Structure of carfilzomib. (D) Dose-response curve of carfilzomib-induced inhibition of *P. vivax* schizont growth area (yellow, IC50 288nM), hypnozoite quantity per well (blue, IC_50_ 511nM), and hepatocyte nuclei quantity (orange, IC_50_ 3.32 µM). Data shown are pooled from three independent experiments, each containing duplicate wells at each concentration. Bars represent ± SEM.

**Figure 3—source data 1**

**Data associated with schizonts versus hypnozoites analyses.** (A) Differentially expressed genes defining schizonts and hypnozoites. Positive Average log 2 FC: Upregulated in hypnozoites relative to schizonts. Rows highlighted in yellow denote inclusion cutoffs for genes considered differentially expressed (Bonferroni adjusted p value < 0.01 & absolute (average log2 fold change [FC]) > 0.5). (B) Differentially expressed genes identified as encoding for cellular fating RNA-binding proteins. (C) GO Term enrichment of Hypnozoite or Schizont transcripts identified as cellular fating RBPs. Rows highlighted in light green are terms with Bonferroni adjusted p value < 0.05. Input parameters: P value cutoff = 0.05; Limit to GO Slim terms = No; Evidence: Computed, curated; PlasmoDB v55.

**Figure 4—supplement 1**

**Assessment of hypnozoite transcriptomes.** (A) UMAP of hypnozoites colored by cluster and faceted by day post-infection. (B) Violin plot showing the average log_2_FC of differentially expressed genes in each cluster. Markers were identified using Seurat’s AllFindMarkers function. Genes with an Bonferroni adjusted p value < 0.05 (Wilcoxon rank-sum) and average log_2_FC > 0.5 were deemed markers. Red dashed line: average log_2_FC = 0.5.

**Figure 4—source data 1**

**Data associated with analyses of hypnozoite-to-hypnozoite heterogeneity.** (A) Markers defining each hypnozoite cluster. Marker defined as: Average log 2 FC > 0.5, Adjusted p value < 0.05. Rows colored green are transcripts that meet marker criteria. (B) Differentially expressed genes identified as encoding for cellular fating RNA-binding proteins in hypnozoite clusters. (C) GO Term enrichment of hypnozoite transcripts identified as cellular fating RBPs. Rows highlighted in light green are terms with Bonferroni adjusted p value < 0.05. P value cutoff = 0.05; Limit to GO Slim terms = No; Evidence: Computed, curated; PlasmoDB v55.

**Figure 5—supplement 1**

**Analysis of host-pathogen transcriptional signatures in *P. vivax-*infected and non-infected hepatocytes.** (A) Number of hepatocytes assessed. Colored by infection status and faceted by sample (left); proportion of infected, exposed, or naive hepatocytes assessed on day 5 or day 9 post-infection. (B) t-SNE plot of hepatocytes colored by day. (C) t-SNE plots of hepatocytes colored by percent expression of zonation markers in human hepatocytes. Markers obtained from Macparland et al. (2019). (D) t-SNE plots of hepatocytes colored by hypnozoite infection status. Grey, uninfected (naive and exposed) hepatocytes; red, hepatocytes infected with C1 hypnozoites; blue, hepatocytes infected with C2; teal, hepatocytes infected with C3 hypnozoites. (E) Unique and overlapping gene sets in hepatocytes infected with schizonts or hypnozoites. Positive enrichment derived from genes displaying greater expression versus non-infected hepatocytes; negative enrichment derived from transcripts displaying decreased expression versus noninfected hepatocytes.

**Figure 6—figure supplement 1**

**Human AKR1B10 is upregulated in hepatocytes infected with *P. vivax***. (A) Violin plots displaying the distribution of *AKR1B10* transcript levels in infected (schizonts and hypnozoites) and non-infected (exposed and naive) hepatocytes. Black dot: mean expression (log_2_). Adj p: Bonferroni adjusted p value versus non-infected (naïve and exposed) hepatocytes. (B) Representative confocal image from a second biological replicate of *P. vivax* parasites on day 12 post-infection of hepatocytes. Cells were stained with DAPI (blue), *Pv*UIS4 (green), and *Hs*AKR1B10 (red). White bar represents 100µm. (D) Associated with Figure 6E (right), violins plots displaying the distribution of AKR1B10 signal across all biological replicates. IFA: immunofluorescence assay; SZ: hepatocyte containing schizont; HZ: hepatocyte containing hypnozoite; EX: hepatocyte not infected but exposed to *P. vivax.* ****: p <= 0.0001; *: p <= 0.05; n.s: not significant, p > 0.05.

**Figure 6—source data 1**

**Data associated with host-specific transcriptional signatures in response to *P. vivax* infection.** (A) Differentially expressed genes in hepatocytes containing schizonts versus non-infected cells. Positive average log2FC: greater expression in infected hepatocytes. (B) Differentially expressed genes in hepatocytes containing hypnozoites versus non-infected cells. Positive average log2FC: greater expression in infected hepatocytes. (C) Enrichment of GO biological processes and Reactome metabolic pathways. (D) Unique and overlapping enriched gene sets in hepatocytes infected with schizonts or hypnozoites. (E) Differentially expressed genes in non-infected hepatocytes exposed to *P. vivax* (exposed) versus non-infected hepatocytes never exposed to *P. vivax* (naïve). Postive average log2FC: greater expression in non-infected hepatocytes exposed to *P. vivax* (exposed). Seurat FindMarkers parameters used to generate gene list: logfc.threshold = 0.25, min.pct = 0.25, only.pos = F, max.cells.per.ident = 20731, assay = “RNA”. (F) Enrichment of GO biological processes and REACTOME metabolic pathways that are downregulated in non-infected hepatocytes exposed to *P. vivax* (exposed) versus non-infected hepatocytes never exposed to *P. vivax* (naïve). Genes in **Figure 6— source data 1E** highlighted in green were used as input to generate enrichment lists.

**Figure 7—source data 1**

**Transcription factor enrichment analysis.** Rows highlighted in green are sets that are significantly enriched, adjusted p value < 0.05.

## Notes

### Competing Interest Statement

The authors have declared no competing interest.

### Summary of Updates

Manuscript text updated. Figures updated. Supplemental files updated.

