## Supplementary figures and images for "Single-cell RNA profiling of *Plasmodium vivax*-infected hepatocytes reveals parasite- and host- specific transcriptomic signatures and therapeutic targets"

### Figure 1 figure supplement 1

#### Figure 1—figure supplement 1

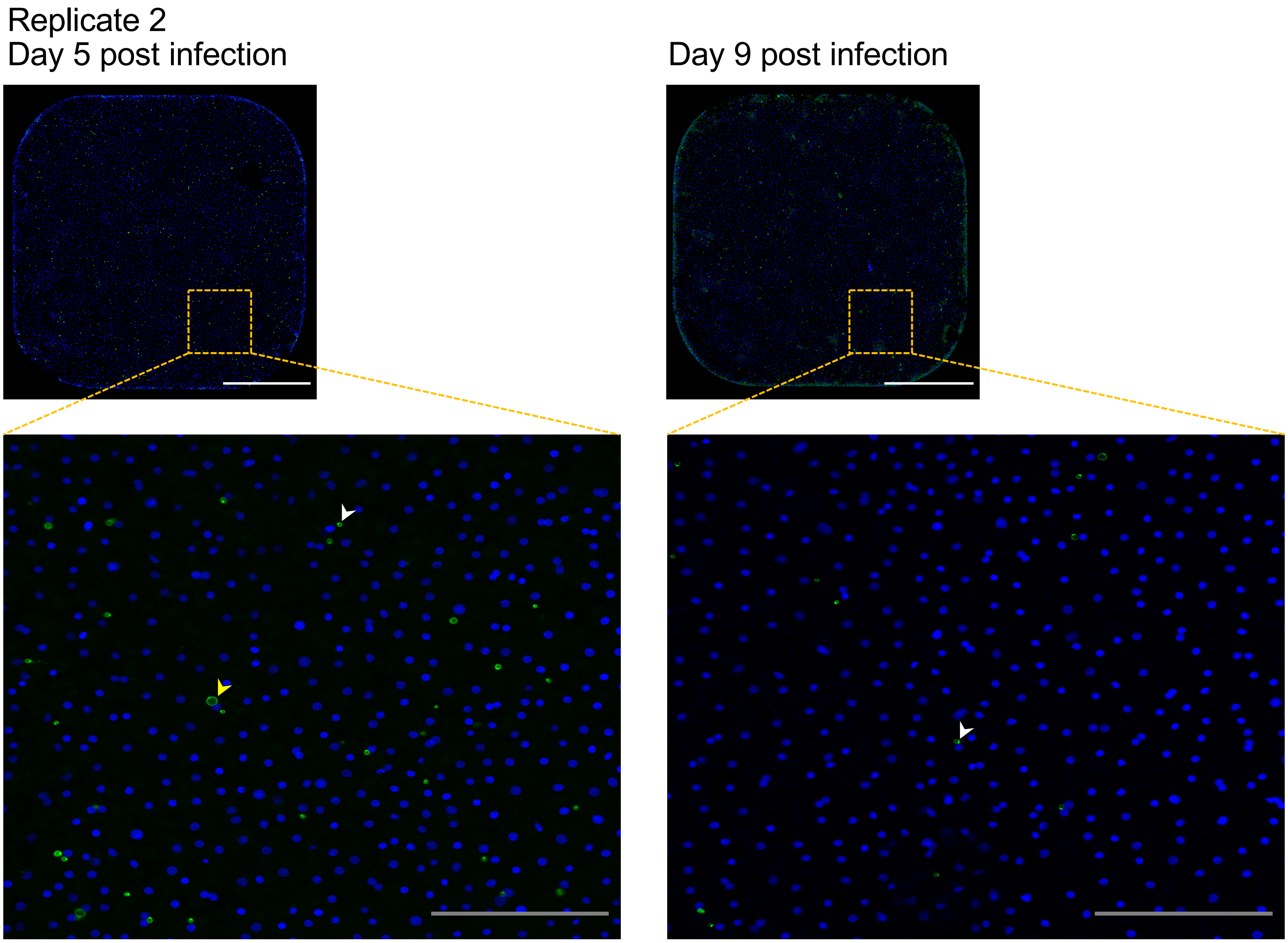

### Figure 2 figure supplement 1

#### Figure 2—figure supplement 1

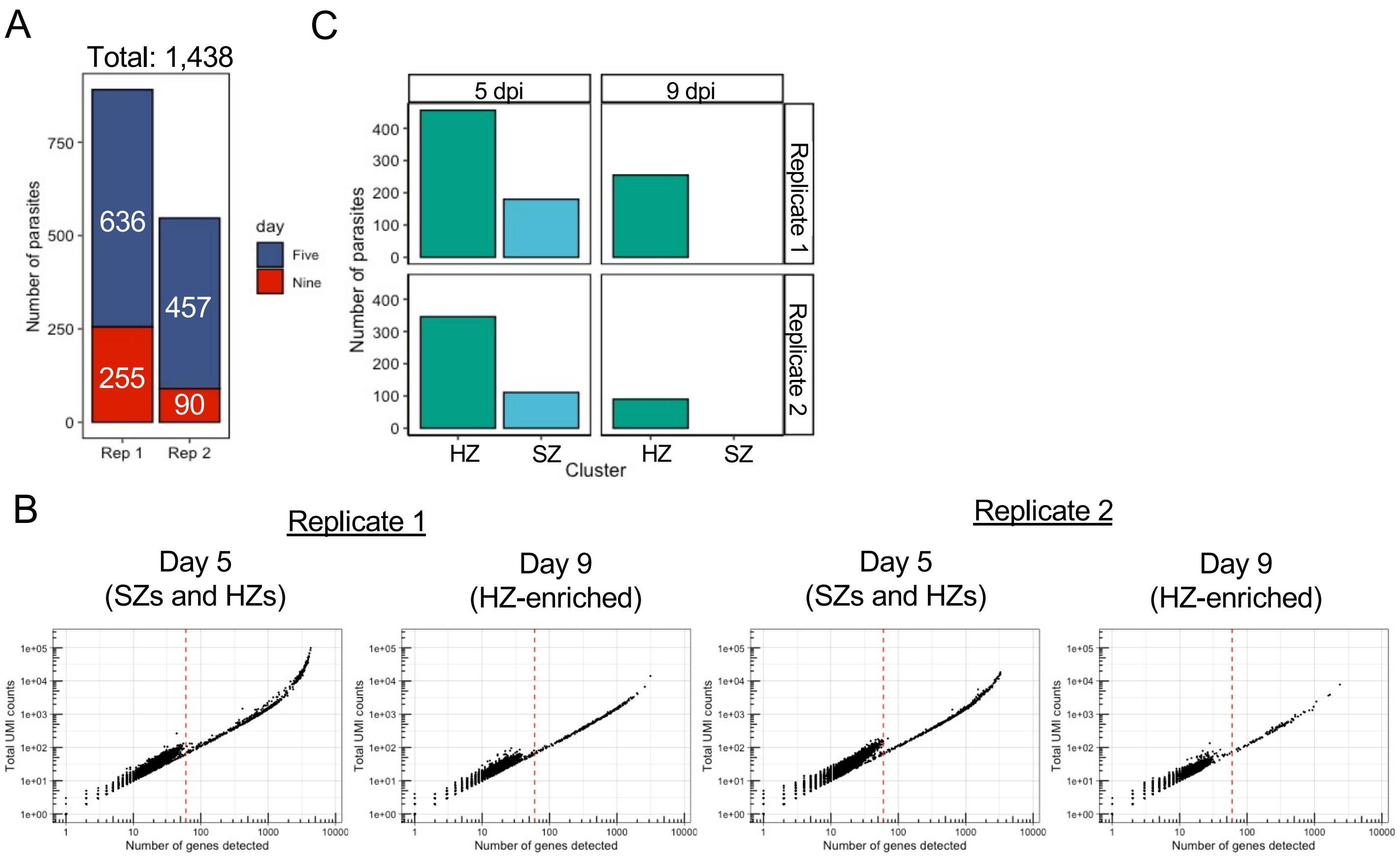

### Figure 2 figure supplement 2

#### Figure 2—figure supplement 2

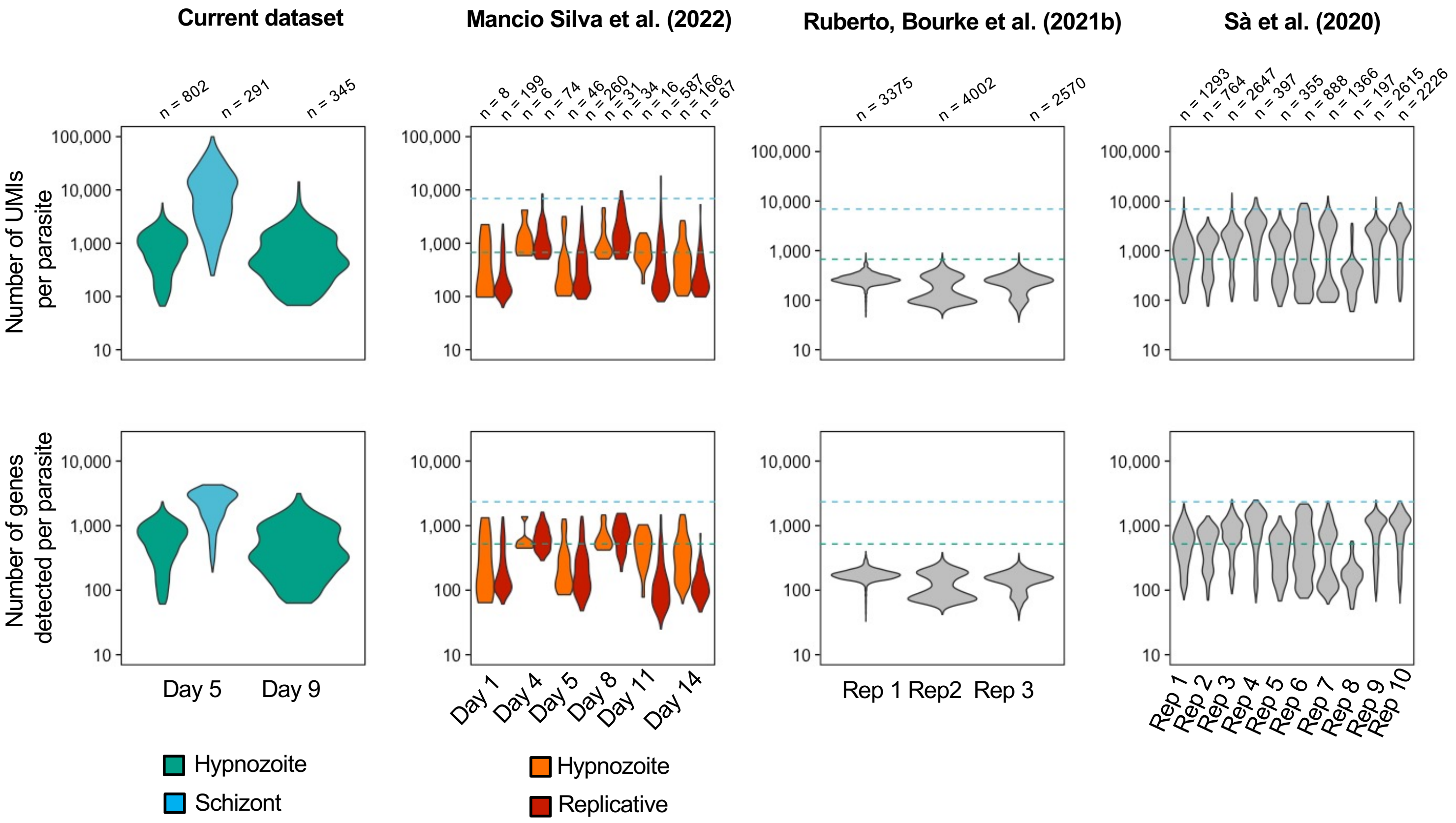

### Figure 3 figure supplement 1

#### Figure 3—figure supplement 1

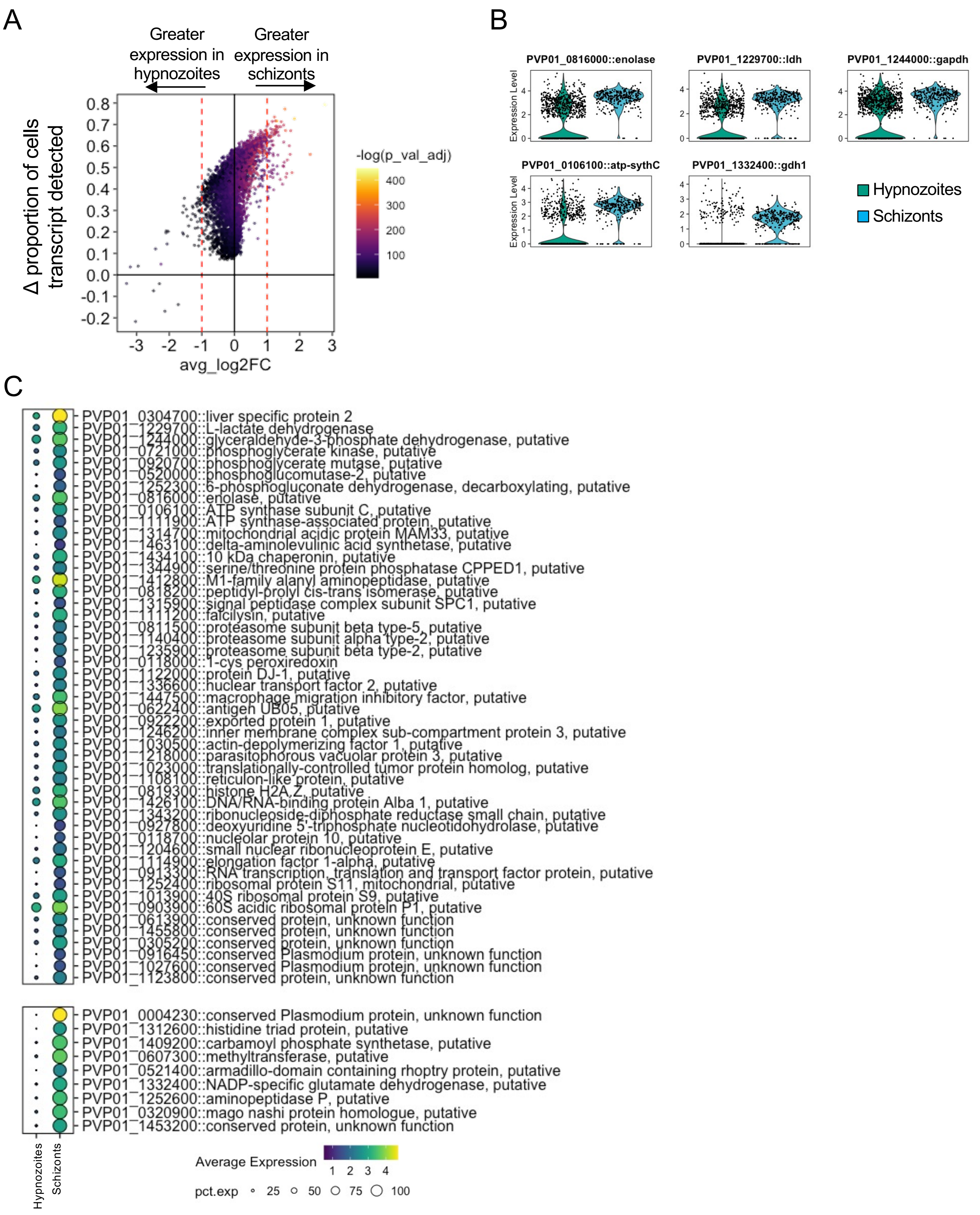

### Figure 4 figure supplement 1

#### Figure 4—figure supplement 1

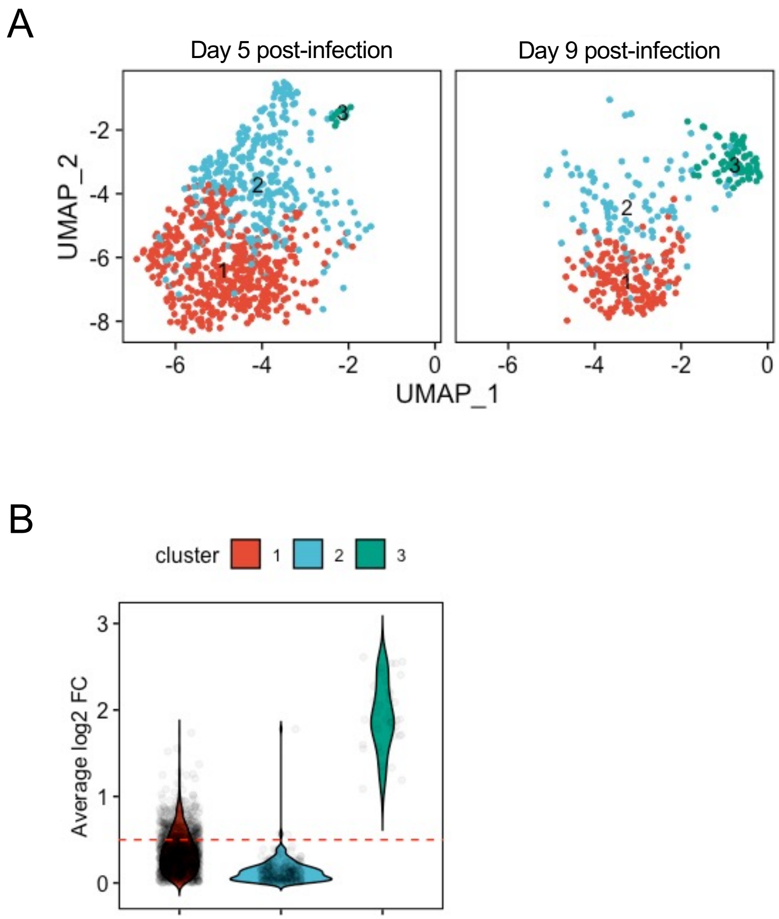

### Figure 5 figure supplement 1

#### Figure 5—figure supplement 1

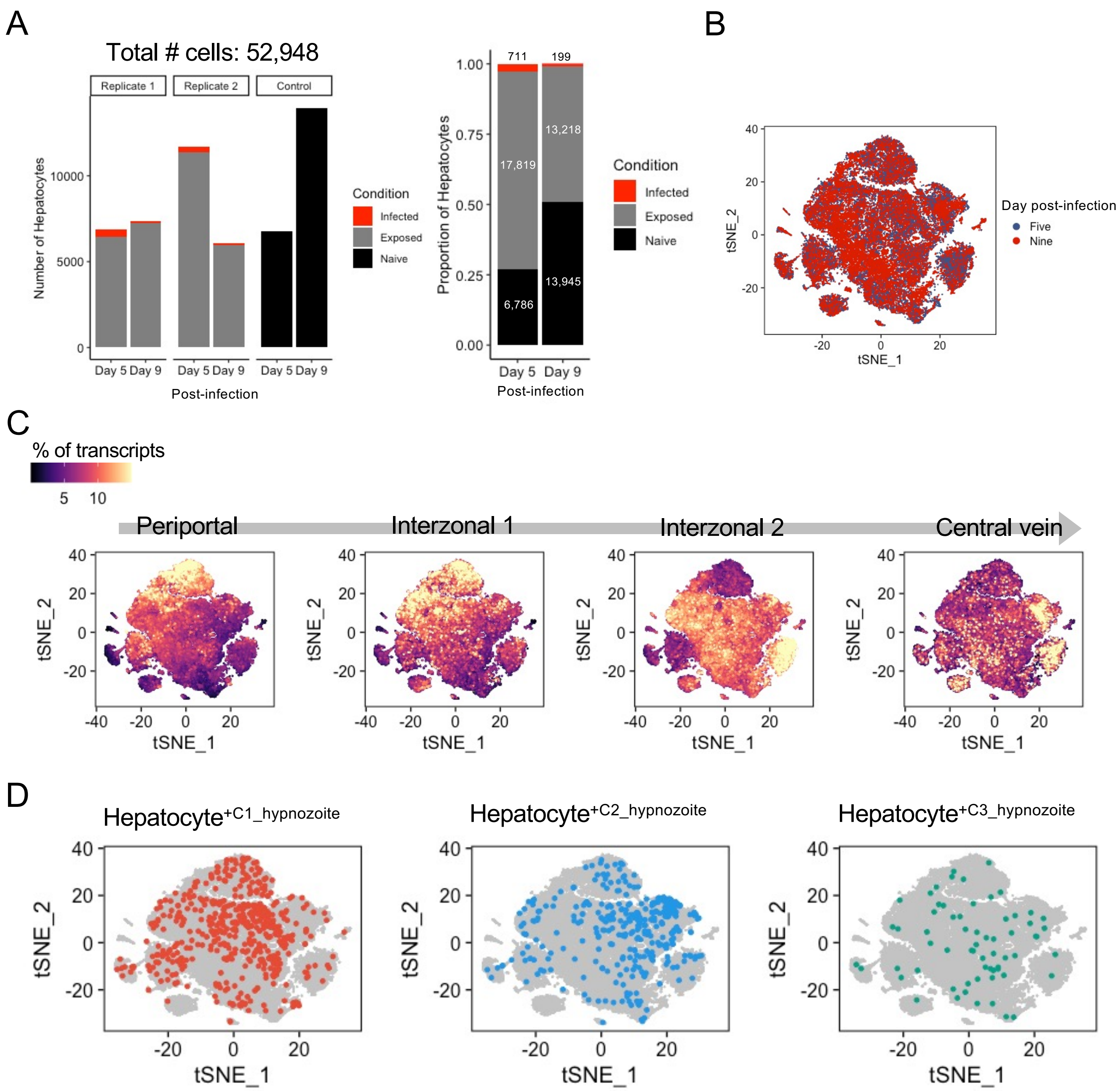

### Figure 6 figure supplement 1

#### Figure 6—figure supplement 1

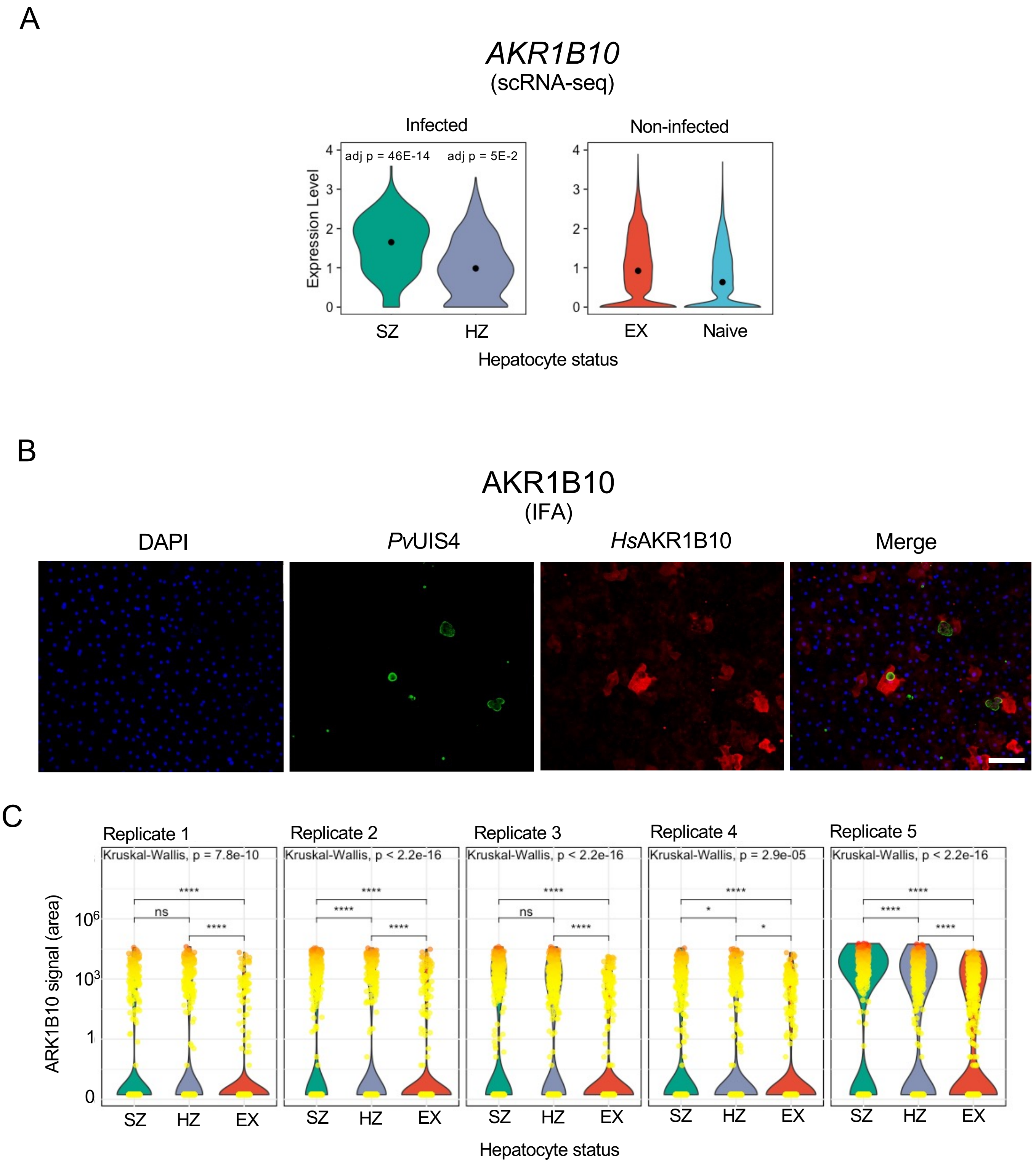
