## Supplementary material for "Single-cell RNA profiling of *Plasmodium vivax*-infected hepatocytes reveals parasite- and host- specific transcriptomic signatures and therapeutic targets": Figure 3 figure supplement 2

A

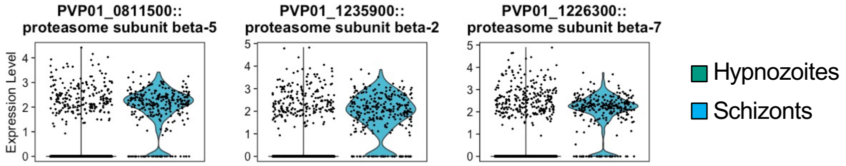


B

| Gene::Description | Average log2FC | Proportion of schizonts transcript detected | Proportion of hypnozoites transcript detected | Adj P Value |
|---|---|---|---|---|
| PVP01_0811500::proteasome subunit beta type-5 | 1.417 | 0.883 | 0.194 | 2.50E-94 |
| PVP01_1140400::proteasome subunit alpha type-2 | 1.356 | 0.856 | 0.207 | 5.39E-81 |
| PVP01_1235900::proteasome subunit beta type-2 | 1.285 | 0.856 | 0.191 | 1.62E-84 |
| PVP01_1226300::proteasome subunit beta type-7 | 0.967 | 0.863 | 0.229 | 3.24E-68 |
| PVP01_1117800::proteasome subunit alpha type-7 | 0.838 | 0.79 | 0.201 | 2.98E-56 |
| PVP01_1448600::ATP-dependent protease subunit ClpQ | 0.823 | 0.814 | 0.179 | 6.28E-68 |
| PVP01_0929900::26S proteasome regulatory subunit RPN7 | 0.763 | 0.856 | 0.225 | 2.36E-57 |
| PVP01_1005800::proteasome maturation factor UMP1 | 0.712 | 0.653 | 0.073 | 2.06E-92 |
| PVP01_1269700::proteasome subunit alpha type-1 | 0.696 | 0.859 | 0.229 | 2.89E-59 |
| PVP01_0206900::proteasome subunit beta type-3 | 0.676 | 0.845 | 0.267 | 3.45E-48 |
| PVP01_1117900::proteasome subunit alpha type-4 | 0.643 | 0.849 | 0.252 | 1.43E-50 |
| PVP01_0112600::26S proteasome regulatory subunit RPN10 | 0.567 | 0.574 | 0.076 | 7.61E-71 |
| PVP01_1015600::proteasome subunit beta type-1 | 0.565 | 0.866 | 0.312 | 1.92E-41 |
| PVP01_0823100::proteasome subunit alpha type-3 | 0.553 | 0.828 | 0.211 | 3.43E-54 |
| PVP01_0822300::26S proteasome non-ATPase regulatory subunit 9 | 0.550 | 0.739 | 0.142 | 5.95E-65 |
| PVP01_1412500::26S protease regulatory subunit 7 | 0.511 | 0.818 | 0.228 | 7.56E-47 |

C

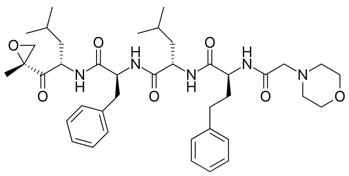

D

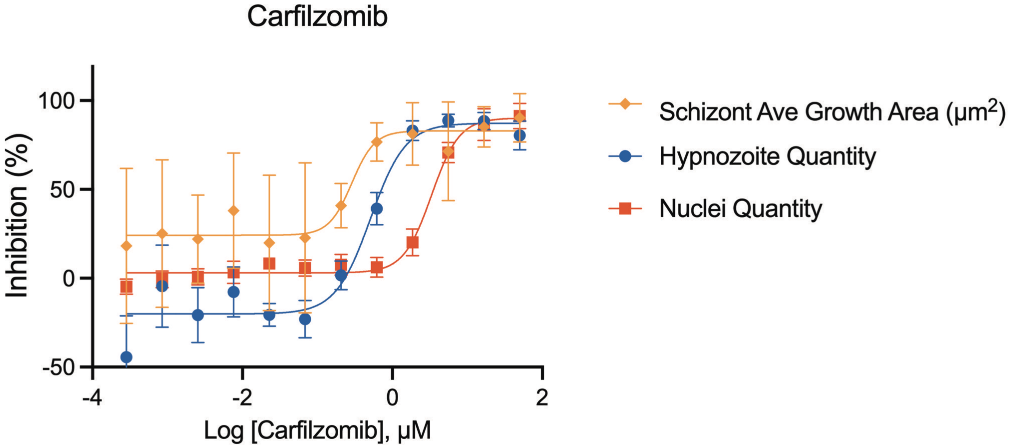
